# Hsp70 chaperones, Ssa1 and Ssa2, limit poly(A) binding protein aggregation

**DOI:** 10.1101/2025.01.17.633617

**Authors:** Hannah E. Buchholz, Sean A. Martin, Jane E. Dorweiler, Claire M. Radtke, Adam S. Knier, Natalia B. Beans, Anita L. Manogaran

## Abstract

Molecular chaperones play a central role in maintaining protein homeostasis. The highly conserved Hsp70 family of chaperones have major functions in folding of nascent peptides, protein refolding, and protein aggregate disassembly. In yeast, loss of two Hsp70 proteins, Ssa1 and Ssa2, is associated with decreased cellular growth and shortened lifespan. While heterologous or mutant temperature sensitive proteins form anomalous large cytoplasmic inclusions in *ssa1Δssa2Δ* strains, it is unclear how endogenous wildtype proteins behave and are regulated in the presence of limiting Hsp70s. Using the wildtype yeast Poly A binding protein (Pab1), which is involved in mRNA binding and forms stress granules (SGs) upon heat shock, Pab1 forms large inclusions in approximately half of *ssa1Δssa2Δ* cells in the absence of stress. Overexpression of Ssa1, Hsp104, and Sis1 almost completely limits the formation of these large inclusions in *ssa1Δssa2Δ*, suggesting that excess Ssa1, Hsp104 and Sis1 can each compensate for the lower levels of Ssa proteins. Upon heat shock, SGs also form in cells whether large Pab1 inclusions are present or not. Surprisingly, cells containing only SGs disassemble faster than wildtype, whereas cells with both large inclusions disassemble slower albeit completely. We suspect that disassembly of these large inclusions is linked to the elevated heat shock response and elevated Hsp104 and Sis1 levels in *ssa1Δssa2Δ* strains. We also observed that wildtype cultures grown to saturation also form large Pab1-GFP inclusions. These inclusions can be partially rescued by overexpression of Ssa1. Taken together, our data suggests that Hsp70 not only plays a role in limiting unwanted protein aggregation in normal cells, but as cells age, the depletion of active Hsp70 possibly underlies the age-related aggregation of endogenous proteins.

## INTRODUCTION

Transient stress such as heat shock can have detrimental impacts to the cell. Cells respond to these stressors by activating protective measures to promote survival such as the heat shock response (HSR), which stimulates molecular chaperone expression. Importantly, these molecular chaperones maintain proteome fidelity by refolding proteins, targeting proteins for degradation, limiting protein aggregation, and regulating stress response pathways (reviewed in Gidalevitz et al., 2011; Lindquist, 1986; Pessa et al., 2024). While much is known about the chaperone response to stress, how chaperones limit unwanted protein aggregation under normal conditions is poorly understood.

The yeast *Saccharomyces cerevisiae* has greatly advanced our understanding of the role of molecular chaperones in both non-stress and stress conditions (Chernova et al., 2017; Ruger-Herreros et al., 2024). Yeast have conserved chaperone families such as J-domain proteins (JDP; otherwise known as Hsp40s), Hsp70s, Hsp90s, and small heat shock proteins, as well as the unique fungal disaggregase, Hsp104. These chaperones work together to maintain protein homeostasis (Chernova et al., 2017; Verghese et al., 2012). Specifically, Hsp70 members mediate the disassembly of protein aggregates, and regulate the HSR, making them one of the most versatile of the chaperone families (Clerico et al., 2015).

Hsp70s in yeast was first identified by Craig and co-workers over 40 years ago (Craig and Jacobsen, 1984). Four members within the Hsp70 family, Ssa1-4, represent major Hsp70 cytosolic proteins in yeast. Knockout of all four isoforms is lethal, whereas presence of at least one isoform is sufficient for viability (Werner-Washburne et al., 1987). Ssa1 and Ssa2 are constitutively expressed, while Ssa3 and Ssa4 are stress-inducible (Werner-Washburne et al., 1987). Despite high sequence similarity, Ssa1-4 isoforms are not necessarily functionally redundant (Young and Craig, 1993). For example, Ssa1 and Ssa3 can maintain the yeast prion [*PSI^+^*] but the other isoforms cannot (Hasin et al., 2014). Furthermore, Ssa4 overexpression does not compensate for the recruitment of Hsp104 to aggregates in *ssa1Δssa2Δ* cells (Andersson et al., 2021), indicating that Ssa4 cannot simply replace all of the functions of the other Ssa proteins. While Ssa proteins are not redundant in all cases, loss of Ssa1 alone appears to have no effect on fitness, whereas loss of both Ssa1 and Ssa2 results in temperature sensitivity, diminished proteostasis, and decreased lifespan (Andersson et al., 2021; Craig and Jacobsen, 1984; Oling et al., 2014), emphasizing the link between these chaperones, stress, and protein aggregation.

Hsp70s also modulate the HSR by inhibiting the major heat shock transcription factor (Hsf1) under non-stress conditions. Ssa1 binds monomeric Hsf1 in the cytoplasm (Zheng et al., 2016), resulting in inhibition of Hsf1 activity (Voellmy and Boellmann, 2007). During transient heat stress, translation machinery, mRNA, and RNA-binding proteins form stress granules (SGs) and Ssa1 is recruited away from Hsf1 (Masser et al., 2019). As a result, unbound Hsf1 enters the nucleus, trimerizes, and binds to heat shock elements (HSEs) to upregulate chaperone gene expression, such as Hsp104 (Anckar and Sistonen, 2011; Gross et al., 1990; Sorger and Nelson, 1989; Voellmy and Boellmann, 2007; Zheng et al., 2016). After stress subsides, chaperones play an important role in the disassembly of SGs. JDP members recognize SG proteins, recruit Hsp70s, and stimulate their ATPase activity. Hsp70s can further recruit Hsp104 to the aggregate, which is responsible for partially threading monomeric substrates from the aggregate through the Hsp104 central pore, resulting in aggregate fragmentation or disassembly (Buchholz et al., 2024; Glover and Lindquist, 1998; Shorter and Lindquist, 2008; Sweeny et al., 2015; Winkler et al., 2012; Yoo et al., 2022). Following SG disassembly, substrate-free Ssa1 can re-bind to Hsf1, inhibit its activity, and ultimately turn off HSR in a negative-feedback loop (Krakowiak et al., 2018; Zheng et al., 2016). The link between Hsp70 and HSR highlights the importance of considering this relationship when studying the influence of Hsp70 on protein aggregates.

Constitutive Ssa members, Ssa1 and Ssa2, appear to limit model proteins from forming protein inclusions under normal conditions. Model proteins, such as the heterologous temperature sensitive NES-LuciTs protein, or mutant yeast Ubc9Ts, *guk1-7*, and *gus1-3* proteins (Andersson et al., 2021; Comyn et al., 2016; Escusa-Toret et al., 2013; Jawed et al., 2022; Jones et al., 2010; Oling et al., 2014; Rolli et al., 2024; Shiber et al., 2013) form inclusions in *ssa1Δssa2Δ* strains. While these studies provide important directions for understanding the role of Hsp70s in limiting protein aggregation, these model proteins are engineered to be temperature sensitive. Therefore, it is unclear how Hsp70s influence proteins that naturally aggregate in response to stress, rather than a byproduct of a mutation. Here, we use the wildtype poly(A) binding protein (Pab1) to study how the loss of major Hsp70 members influence protein states. Under normal conditions, Pab1 is localized to the cytoplasm and binds mRNAs to ensure mRNA stability and promote translation (Brambilla et al., 2019). Under transient heat shock conditions, Pab1 and associated RNAs form SGs (Hoyle et al., 2007; Swisher and Parker, 2010). We find that in *ssa1Δssa2Δ*, Pab1-GFP forms large cytoplasmic inclusions without heat stress. Formation of these inclusions can be limited by the overexpression of Ssa1, the JDP Sis1, or Hsp104, suggesting that any one of these chaperones can override the aberrant aggregation phenotype caused by the lack of Ssa1 and Ssa2. While *ssa1Δssa2Δ* cells can form SGs upon heat shock, the SG puncta in the double mutant resolve faster than SG puncta in wildtype cells. We suspect that the faster recovery is due to an elevated HSR in the double mutant. Interestingly, we show that cells from saturated yeast cultures also form Pab1-GFP large inclusions, which can be partially inhibited by the overexpression of Ssa1. Taken together, we suspect that Hsp70s plays an important role in limiting unwanted protein aggregation in both normal and aging cells.

## RESULTS

### Pab1-GFP forms a large inclusion in ssa1Δssa2Δ cells

To understand the role of Hsp70 in maintaining SG proteins in the absence of stress, we engineered a *SSA1* deletion (*ssa1Δ*) in the 74D-694 genetic background. Using GFP fused to the C-terminus of the poly(A) binding protein, Pab1 (Pab1-GFP), under the control of the *PAB1* promoter (Brengues and Parker, 2007). We found that Pab1-GFP in *ssa1Δ* exhibits diffuse cytoplasmic fluorescence in the absence of stress (or no heat shock) conditions, similar to wildtype strains (Figure 1A; Figure S1). After transient heat shock of 42°C for 45 minutes, Pab1-GFP forms characteristic SGs in wildtype and *ssa1Δ* strains (Figure 1A), revealing loss of Ssa1 alone has no impact on the formation of Pab1 SGs, likely due to the other constitutively expressed isoform of Hsp70, Ssa2 (Werner-Washburne et al., 1987). Therefore, we removed both constitutive Ssa isoforms, *ssa1Δssa2Δ,* to investigate Pab1-GFP aggregate morphology during room temperature and stressed conditions. In comparison to wildtype and *ssa1Δ* strains in the absence of temperature stress, Pab1-GFP forms a single, large inclusion in approximately 20% of *ssa1Δssa2Δ* cells (Figure 1A; Figure S1). Upon heat stress, we made two observations: 1) all *ssa1Δssa2Δ* cells formed small Pab1-GFP puncta, which are consistent with SGs, and 2) the number of *ssa1Δssa2Δ* cells containing large inclusions doubled to an average of 43.7% (Figure 1A). These data suggest that Ssa1 and Ssa2 together do not play a role in SG formation as suggested previously (Walters et al., 2015), but play a role in limiting Pab1 from forming large inclusions in wildtype cells under normal conditions.

**Figure 1.**
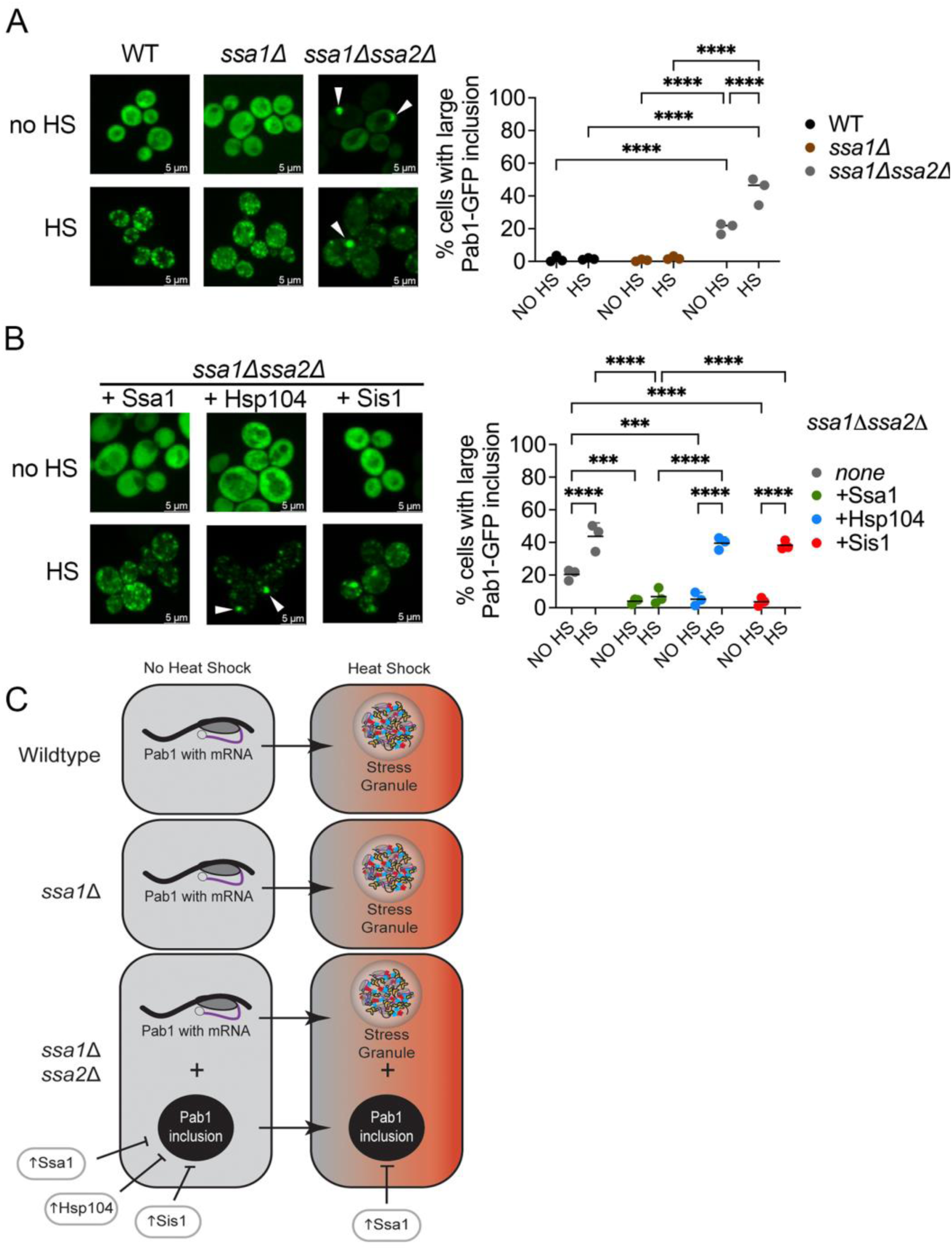
Pab1-GFP forms large inclusions in *ssa1Δssa2Δ* cells, which can be limited by chaperone overexpression. A) Wildtype, *ssa1Δ*, and *ssa1Δssa2Δ* strains containing Pab1-GFP were grown to late log phase. Paired strains were with incubated at room temperature (no heat shock; no HS) or subjected to a 42°C for 45-minute heat shock (HS). Field images were taken using wide-field fluorescent microscopy, and a minimum of 200 cells per replicate were scored for the presence of at least one large Pab1-GFP inclusion. B) Similar to A, the percentage of cells with a large Pab1-GFP inclusion was quantified in a *ssa1Δssa2Δ* strain containing either a GPD-Ssa1, HSE-Hsp104, or GPD-Sis1 overexpression plasmid. For easier comparison, the *ssa1Δssa2Δ* data without plasmid overexpression is the same as shown in A. A two-way ANOVA with Tukey’s multiple comparisons test was used to compare strains in A and B (Only significant comparisons are shown; ***p≤0.0003; ****p≤0.0001). C) Summary of results from figure 1. In the absence of stress, Pab1-GFP (indicated as a protein bound to mRNA in the tan boxes) exhibits diffuse fluorescence, and forms normal stress granules following heat shock in wildtype and *ssa1Δ* cells (shown in the orange boxes to the right). Pab1-GFP forms large inclusions in *ssa1Δssa2Δ* cells in the absence of stress, which can be limited by Ssa1, Hsp104, or Sis1 overexpression. However, only Ssa1 can limit Pab1-GFP inclusions following heat shock in *ssa1Δssa2Δ* cells. Medians are shown as horizontal lines.

To confirm whether the large Pab1-GFP inclusion observed in Figure 1A is dependent on Hsp70, we introduced a plasmid that constitutively expresses *SSA1* under control of a *GPD* promoter (GPD-Ssa1) into *ssa1Δssa2Δ* cells. Compared to *ssa1Δssa2Δ* controls that showed an average of 20.4% of cells with a large Pab1-GFP inclusion without heat shock, introduction of Ssa1 showed only 3.9% of cells contained large inclusions in these strains (Figure 1B; Figure S1; Figure 1C left). Since both Hsp104 and Sis1 work in concert with Hsp70s, we also asked whether overexpression of these other chaperones would also prevent the formation of large Pab1-GFP inclusions without heat shock. Using a plasmid where *HSP104* expression is controlled by the native *HSP104* promoter (HSE-Hsp104) and increases Hsp104 levels four-fold over presence of the endogenous copy alone (Buchholz et al., 2024; Howard et al., 2020; Jackrel and Shorter, 2014), only 5.2% of cells contained Pab1-GFP inclusions in *ssa1Δssa2Δ* cells, which is similar to wildtype cells under no heat shock conditions. Similarly overexpressing *SIS1*, under the control of a constitutive *GPD* promoter (GPD-Sis1) resulted in 3.6% of cells with large inclusions without heat shock (Figure 1B; Figure S1). These data indicate that under normal conditions, extra Ssa1, Hsp104, or Sis1 can limit the formation of large Pab1-GFP inclusions in *ssa1Δssa2Δ* cells (Figure 1C, left).

We then determined whether Ssa1, Hsp104, or Sis1 influences Pab1-GFP aggregation following heat stress. Similar to non-stress conditions, introduction of GPD-Ssa1 in heat shocked cultures resulted in the formation of SGs in all cells, yet only 6.8% of cells contained large Pab1-GFP inclusions (Figure 1B; Figure S1; Figure 1C, right), suggesting that the presence of Ssa1 is sufficient to limit Pab1-GFP inclusion formation during stress. However, strains containing the HSE-Hsp104 or GPD-Sis1 plasmid formed SGs in all *ssa1Δssa2Δ* cells and on average 39.4 and 38.2% of the cells, respectively, also contained large Pab1-GFP inclusions under stress conditions (Figure 1B, Figure S1, Figure 1C, right). We suspect that these Pab1-GFP inclusions are newly formed since they are not readily detectable under non-heat stressed conditions (Figure 1B). While overexpression of Hsp104 or Sis1 appear to limit large Pab1-GFP inclusions under non-stress conditions, overexpression of either chaperone is insufficient to limit large inclusion formation upon heat stress. In contrast, Ssa1 is sufficient to limit Pab1-GFP inclusion formation under heat shock conditions in *ssa1Δssa2Δ* strains (Figure 1B, Figure 1C, right).

### Sup35 prion domain inclusions in ssa1Δssa2Δ strains colocalize with Pab1-mCherry inclusions

Proteins involved in translation, such as Pab1, Pub1, and Sup35, often contain intrinsically disordered regions (IDRs) and aggregate upon stress (Buchan and Parker, 2009; Franzmann et al., 2018; Wallace et al., 2015). These proteins appear to be associated with different parts of the SG, which consists of a ‘core’ that assembles first through stronger interactions followed a dynamic ‘shell’ that has weaker interactions (Jain et al., 2016; Wheeler et al., 2016). Interestingly, Pab1 is considered a core SG protein, but Sup35 is not (Jain et al., 2016). Therefore, we asked if the IDR of Sup35, specifically the disordered N-terminal and middle domains (called the prion domain, PrD), form inclusions in *ssa1Δssa2Δ* strains in the absence of heat shock.

We transformed two plasmids into wildtype and *ssa1Δssa2Δ* strains: one plasmid containing Sup35 PrD fused to GFP (PrD-GFP) under a copper inducible (pCUP1) promoter, and another plasmid containing Pab1 with a C-terminal mCherry tag (Pab1-mCherry) under the *PAB1* promoter. In wildtype strains, both PrD-GFP and Pab1-mCherry exhibit diffuse fluorescence under no heat shock conditions. However, only Pab1-mCherry forms small puncta like SGs during heat stress (Figure 2A, Figure S2). In *ssa1Δssa2Δ* cells, PrD-GFP forms large inclusions in 6-8% of cells, regardless of heat shock or not (Figure 2A). However, in the small number of cells that contained PrD-GFP inclusions under no heat shock, most of these inclusions colocalized with the large Pab1-mCherry inclusions (Figure 2A; Figure S2). During heat shock, we would see an occasional colocalization between PrD-GFP and Pab1-mCherry (Figure 2A), but most of the time we would see no overlap (Figure S2). It is unclear which regions of Sup35 are responsible for biologically relevant phase separation *in vivo* upon heat shock. It has been suggested that the disordered regions of the PrD can phase separate (Franzmann et al., 2018; Grizel et al., 2024). However others argue the essential C-terminal GTPase domain, which is required for translation and not included in our constructs, allow for phase separation *in vivo* (Grimes et al., 2023). Our data suggest that the inclusions that form without temperature stress could also contain other IDR proteins. However, it is unclear whether these large inclusions associated with temperature stress have similar composition to inclusions present in non-stressed conditions.

**Figure 2.**
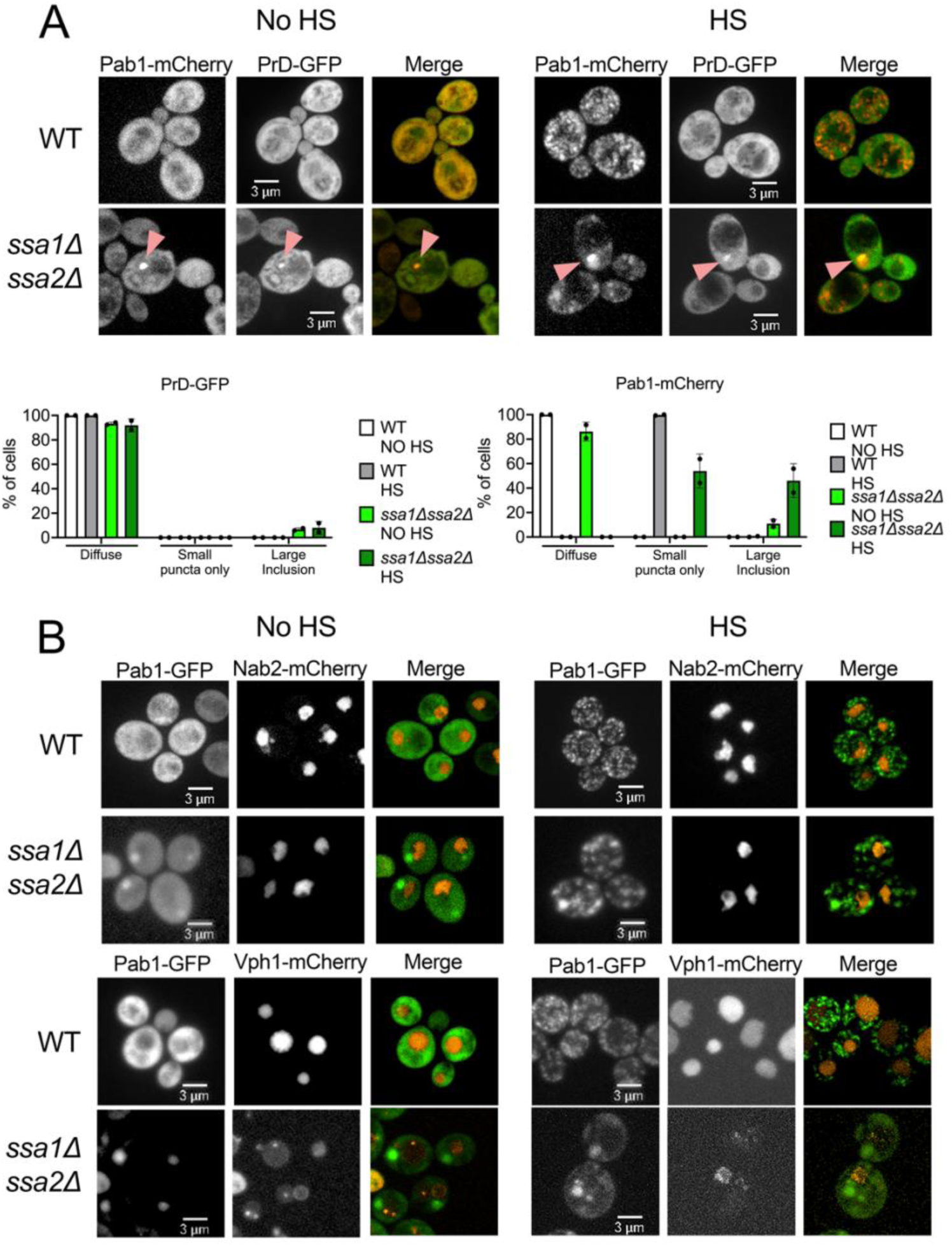
Pab1-GFP inclusions co-localize with prion domain of Sup35. A) *Top* The indicated strains were transformed with Pab1-mCherry, under the *PAB1*promoter, and the prion domain of Sup35 (PrD-GFP) under the copper inducible *CUP1* promoter. Cells were grown overnight in the presence of 25 µM CuSO_4_.to induce PrD-GFP expression. Cells were either left at room temperature (no HS) or heat shocked (HS) and visualized with spinning disk confocal microscopy. Pink arrows indicate colocalization. Other trials can be found in Supplemental Figure 2. *Bottom* The percentage of cells exhibiting PrD-GFP (left) or Pab1-mCherry (right) diffuse fluorescence, only small puncta (stress granules), or large inclusions were quantified from acquired images. At least 30 cells of two trials were analyzed for each condition. B) Wildtype and *ssa1Δssa2Δ* strains were integrated with either Nab2-mCherry to visualize the nucleus (top), or Vph1-mCherry to visualize the vacuole (bottom), and transformed with Pab1-GFP. Cells were visualized either with heat treatment (left) or without heat shock (right) with spinning disk confocal microscopy. Other trials can be found in Supplemental Figure 2.

We next asked whether the large Pab1-GFP inclusions are located specifically near the nucleus or the vacuole. The temperature-sensitive *guk1-7*-GFP forms perinuclear aggregates in *ssa1Δssa2Δ* cells (Andersson et al., 2021). The large Pab1-GFP inclusions in *ssa1Δssa2Δ* cells did not appear to be perinuclear or co-localize with the nuclear Nab2-mCherry marker in either no heat shock or heat shock conditions (Figure 2B, Figure S2). The insoluble protein deposit (IPOD) is located adjacent to the vacuole and contains terminally aggregated proteins such as the Rnq1 prion and expanded polyQ repeats of Huntingtin (Escusa-Toret et al., 2013; Kaganovich et al., 2008; Rolli et al., 2024). To determine whether Pab1 inclusions in *ssa1Δssa2Δ* were near the vacuole, we used the vacuolar membrane marker, Vph1-mCherry. Interestingly, Vph1 shows clear vacuolar staining in wildtype cells (Figure 2B), yet appears to have dim vacuolar fluorescence in *ssa1Δssa2Δ.* Since the vacuolar uptake of cytosolic proteins is decreased in *ssa1Δssa2Δ* strains (Horst et al., 1999), it is possible that Vph1 localization to the vacuole could also be compromised. Despite the low vacuolar fluorescence, it appears that the *ssa1Δssa2Δ* large inclusions are localized near to the vacuole, rather than perinuclear localization that has been reported for other model proteins in *ssa1Δssa2Δ* strains (Andersson et al., 2021).

### ssa1Δssa2Δ cells exhibit an elevated HSR and increased chaperone levels

As previously discussed, Hsp70 is linked with Hsf1 activity and the HSR. To verify that Hsf1 activity is elevated in our *ssa1*Δ*ssa2*Δ strain, we used a HSE-YFP reporter gene, in which the yellow fluorescent protein is driven by a promoter with four tandem Hsf1 binding motifs (HSEs), and can indirectly measure Hsf1 activity by the level of YFP using flow cytometry (Zheng et al., 2016). We found that, both with and without heat shock, HSE-YFP levels in *ssa1*Δ*ssa2*Δ cells were 14-fold greater compared to wildtype and *ssa1Δ* cells (Figure 3A), verifying that HSR is increased in these mutants.

**Figure 3.**
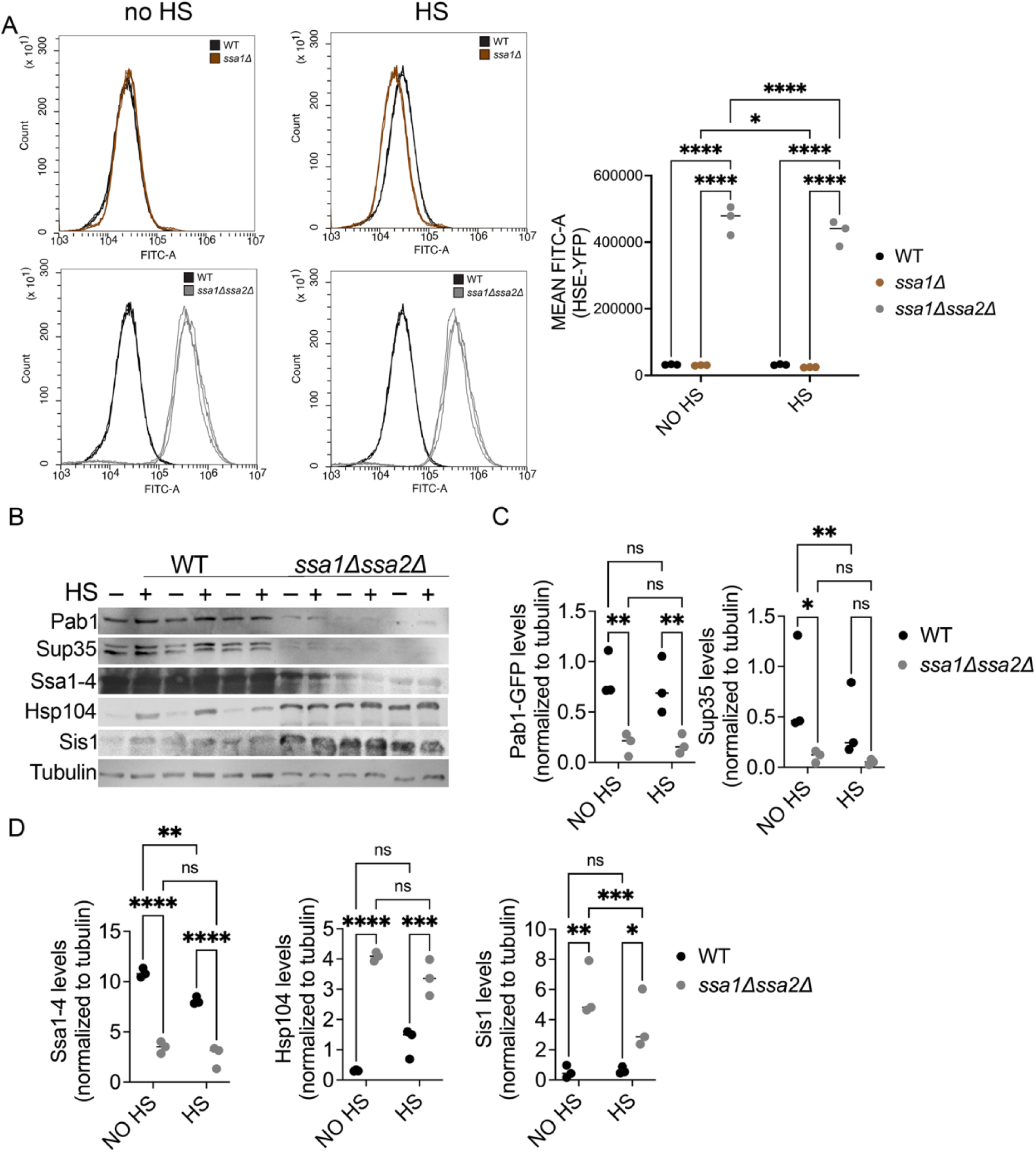
*ssa1Δssa2Δ* strains exhibit an elevated heat shock response and increased chaperone levels. A) WT, *ssa1,* and *ssa1Δssa2Δ* strains were integrated with an HSE-YFP reporter (Zheng et al., 2016). Three independent cultures of each strain were grown to late log and assessed for YFP intensity through flow cytometry using the FITC-A filter without (left, no HS) or following heat shock (middle, HS). Mean FITC area of each individual trial are compared by as indicated (right). B) Three independent paired cultures of WT and *ssa1Δssa2Δ* strains were grown to late log and either heat shocked (HS) at 42°C or incubated at room temperature (no HS) for 45 minutes. Cells were lysed approximately 10 minutes after treatment, lysates were immediately run on SDS-PAGE and subjected to Western blot analysis using the indicated antibodies. C) Steady state levels of translation-associated proteins (Pab1-GFP and Sup35) within and between no HS and HS strains were compared. D) Steady state levels of chaperone proteins (Ssa1-4, Hsp104, and Sis1) within and between no HS and HS strains are shown. All statistical tests involved a two-way ANOVA with an uncorrected Fisher’s LSD test (*p≤0.038; **p≤0.0099; ***p=0.0007; ****p≤0.0001). Medians are indicated with horizontal lines.

Except for certain molecular chaperone genes, global translation is generally repressed during transient heat stress (Anderson and Kedersha, 2006; Kedersha and Anderson, 2002). Therefore, we assessed protein steady state levels of non-chaperone proteins (tubulin, Pab1-GFP and Sup35) as well molecular chaperones (Ssa1-4, Hsp104, and Sis1). Heat shock was induced in wildtype and *ssa1Δssa2Δ* strains, and cultures were immediately lysed for Western blot analysis. General levels of tubulin are comparable between wildtype and *ssa1Δssa2Δ* strains with and without heat shock (Figure 3B; Figure S3). The steady state levels of both Pab1 and Sup35 are significantly lower in *ssa1Δssa2Δ* compared to wildtype strains in the absence of heat stress (Figure 3B and 3C). These data imply that the constitutively elevated HSR in *ssa1Δssa2Δ* could limit the accumulation of certain proteins, possibly those that are associated with translation and SG assembly. It is important to note that while overexpression of aggregation-prone or intrinsically disordered proteins can lead to protein aggregation (Chakrabortee et al., 2016; Zhou et al., 2001), these Pab1-GFP inclusions are observed in *ssa1Δssa2Δ* despite lower steady state levels of Pab1-GFP.

Next, we assessed steady state levels of molecular chaperones. Using an antibody that recognizes Ssa1-4, the levels of Ssa1-4 protein detected in wildtype cells is about 3 times more than in *ssa1Δssa2Δ* cells (Figure 3D). The Ssa1-4 signal we observed in *ssa1Δssa2Δ* strains is likely the stress-inducible Hsp70 isoforms (Ssa3 and Ssa4), since it has been reported that Ssa4 levels increase in strains lacking Ssa1 and Ssa2 (Boorstein and Craig, 1990), and we postulate these cells are under a constant state of elevated HSR (Figure 3A). The steady state levels of Hsp104 and Sis1 were significantly increased by approximately ten-fold in the *ssa1Δssa2Δ* strain, suggesting that the elevated HSR in *ssa1Δssa2Δ* is also upregulating these chaperone genes (Figure 3D).

### Transient heat shock does not impact growth of ssa1Δssa2Δ strains

We asked whether transient heat shock treatment of 42°C for 45 minutes would impact growth of *ssa1Δssa2Δ* strains. Wildtype, *ssa1Δ*, and *ssa1Δssa2Δ* were transformed with Pab1-GFP. *ssa1Δssa2Δ* were also transformed with either an empty vector (EV) or a plasmid that contained either GPD-Ssa1, HSE-Hsp104, or GPD-Sis1 used in figure 1. Wildtype and *ssa1Δ* strains did not exhibit growth defects in the presence or absence of heat shock (Figure S4). However, *ssa1Δssa2Δ* showed a substantial growth defect compared to wildtype and *ssa1Δ* strains, regardless of treatment. These results are consistent with previous reports that *ssa1Δssa2Δ* strains do not grow at 37°C and have slower growth at 30°C (Craig and Jacobsen, 1984). This slow growth phenotype is rescued upon the introduction of Ssa1 (Figure S4), which correlates with the lack of the large Pab1-GFP inclusions in *ssa1Δssa2Δ* with Ssa1 overexpression (Figure 1B). Interestingly, overexpression of Hsp104 or Sis1 was unable to significantly improve growth of *ssa1Δssa2Δ* strains in the presence or absence of heat shock (Figure S4), which again is consistent with the inability of these chaperones to prevent Pab1-GFP inclusion formation in *ssa1Δssa2Δ* upon heat shock (Figure 1B).

### SGs recover faster in ssa1Δssa2Δ cells than wildtype cells

The disassembly of SGs after heat shock has been shown to require several chaperones including Hsp104, Hsp70, and J domain proteins (Walters et al., 2015; Kroschwald et al., 2015; Yoo et al., 2022). Sis1 and Ssa1 are thought to initially interact with the condensate and act as a marker for Hsp104 binding. Hsp104 has been proposed to mediate disassembly through a partial threading mechanism that is regulated by the middle domain of the chaperone (Yoo et al., 2022). Based on what is known about the disassembly of SGs, we asked how Pab1-foci disassembly (both SG and large inclusions) is impacted in *ssa1Δssa2Δ* cells.

Pab1-GFP SGs resolve in the BY4741 genetic background as quickly as two hours (Shattuck et al., 2019). We monitored SG recovery in our 74D-694 background by capturing field images of cells each hour following heat shock and quantifying the percentage of cells with and without visible Pab1-GFP SGs or large inclusions. In wildtype cells, Pab1-GFP SGs form in 100% of the cells immediately following heat shock regardless if a large inclusion is present or not. SGs in wildtype cells resolve over time, with less than 10% of cells containing SGs by five hours (Figure 4A). Compared to wildtype cells, which exhibit an inverse sigmoidal curve starting with a slow lag phase followed by rapid decline of cells containing SGs, *ssa1Δ* strains show an extended lag time of approximately 3 hours before cells exhibit linear disassembly of Pab1-GFP over the next few hours. Approximately 25% of the cells still contain SGs after 5 hours in *ssa1Δ* strains, suggesting that both Ssa1 and Ssa2 have unique contributions for SG disassembly or the total concentration of Ssa1 and Ssa2 is important for disassembly. As discussed above during heat shock, all *ssa1Δssa2Δ* cells form SG and the number of cells that contain large Pab1 inclusions increase (Figure 1A). When monitoring disassembly and the transition from cells with GFP aggregates to diffuse fluorescence, we observe that the kinetics of aggregate loss is faster than the other strains (Figure 4A). In fact, we observe on average 34.1% of *ssa1Δssa2Δ* cells have some type of GFP aggregate after two hours compared to the wildtype or *ssa1Δ* strains (Figure 4A). Since *ssa1Δssa2Δ* exhibits elevated HSR, we suspect that the increased chaperone levels in these strains allow for faster aggregate disassembly.

**Figure 4.**
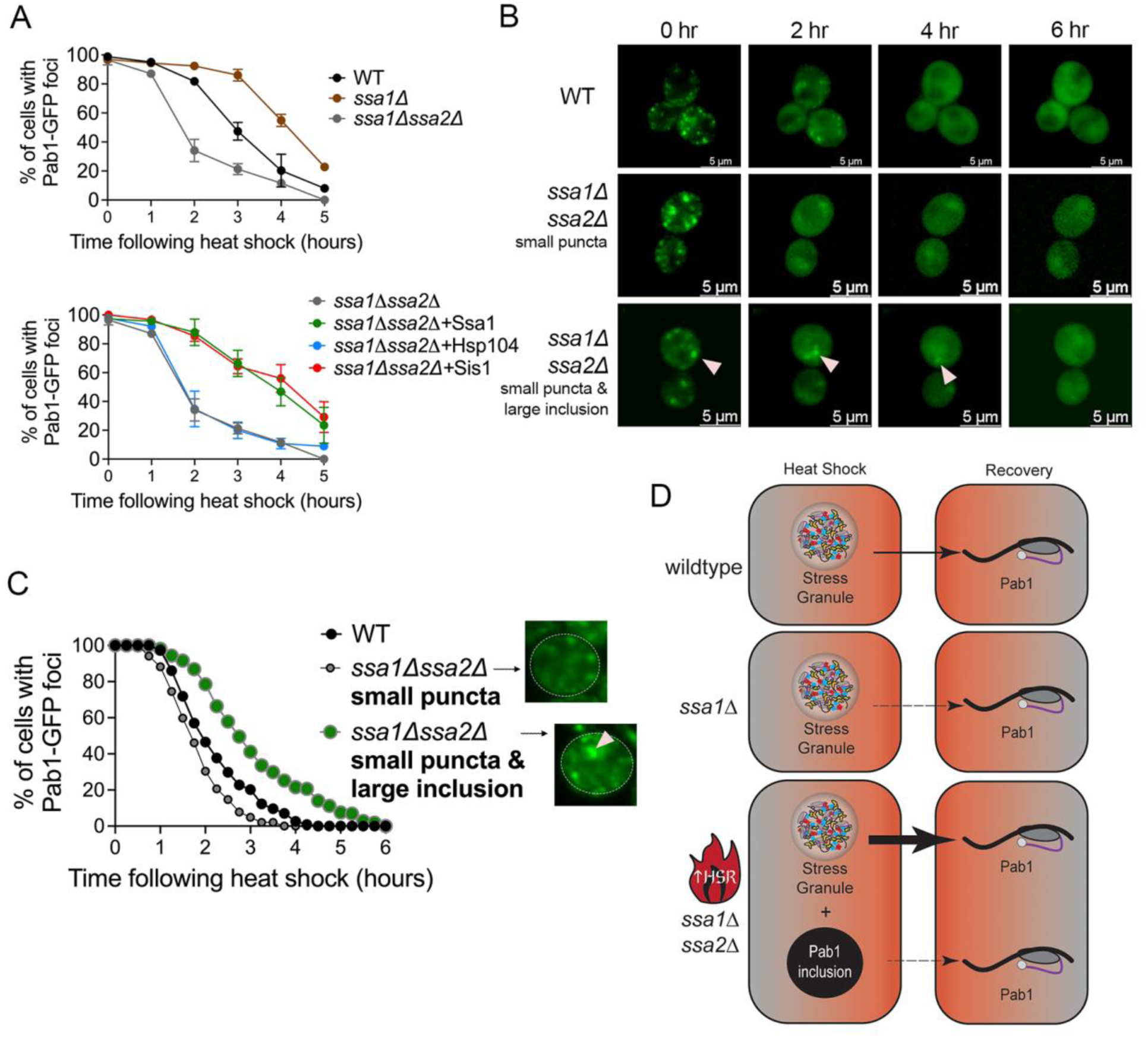
*ssa1Δssa2Δ* cells with large Pab1-GFP inclusions exhibit slower disassembly kinetics compared to cells with only small puncta. A) *Top.* Wildtype, *ssa1Δ*, and *ssa1Δssa2Δ* strains containing Pab1-GFP were heat shocked and fields of cells were imaged for the presence of any Pab1-GFP aggregate (either SG or large inclusion) over five hours. Each data point represents a mean percentage of cells with Pab1-GFP foci from 3-4 independent cultures, with a minimum of 200 cells quantified per time point for each culture. *Bottom.* Pab1-GFP foci disassembly was quantified in *ssa1Δssa2Δ,* similar to the top panel, with the indicated chaperone overexpressed from a plasmid. For easier comparison, the *ssa1Δssa2Δ* data without plasmid overexpression is the same as shown in A. B) Wildtype and *ssa1Δssa2Δ* strains containing Pab1-GFP were heat shocked and subjected to 3D timelapse microscopy, capturing images every 15 minutes. *ssa1Δssa2Δ* cells were categorized by either initially only containing small puncta (middle panel) or initially containing both a large Pab1-GFP inclusion (pink arrow) and small puncta similar to stress granules (bottom panel). Images are shown in two-hour increments. C) At least 100 cells were subjected to 3D-timelapse for each category to monitor the loss of all Pab1-GFP fluorescent aggregates (either cells with puncta or cells containing both puncta and inclusions) in the following cell types: WT cells (which contain only small puncta), *ssa1Δssa2Δ* containing small puncta only (representative image shown at 0 time is shown), or *ssa1Δssa1Δ* containing both small puncta and the large inclusion (representative image is shown, with the arrow showing the large inclusion). The time in all Pab1-GFP foci were lost and the cell exhibited complete diffuse fluorescence was recorded. The graph represents the percentage of cells at any given time point that contain the indicated Pab1-GFP foci. D) Summary results of Figure 4. Pab1-GFP SG recovery is slower in *ssa1Δ* cells compared to wildtype (as indicated by thin arrow). *ssa1Δssa2Δ* cells, with increased HSR, exhibit faster SG recovery compared to wildtype cells (as indicated by a think arrow). *ssa1Δssa2Δ* cells containing a large Pab1-GFP inclusion and SG exhibit slower overall Pab1-GFP recovery.

Overexpression of Ssa1 limits Pab1-GFP inclusion formation in *ssa1Δssa2Δ* cells in the absence and presence of heat shock (Figure 1A and B). Here, we asked whether SG disassembly kinetics would change upon Ssa1 overexpression in *ssa1Δssa2Δ* cells. Introduction of GPD-Ssa1 into *ssa1Δssa2Δ*, which produces a scenario where only Ssa2 is absent, substantially slows down Pab1-GFP foci disassembly (Figure 5B), similar to *ssa1Δ* strains. Therefore, loss of Ssa1 (*ssa1Δ*; Figure 4A Top), or presumably Ssa2 (*ssa1Δssa2Δ* + GPD-Ssa1; Figure 4A Bottom), is insufficient to restore wildtype disassembly kinetics.

**Figure 5.**
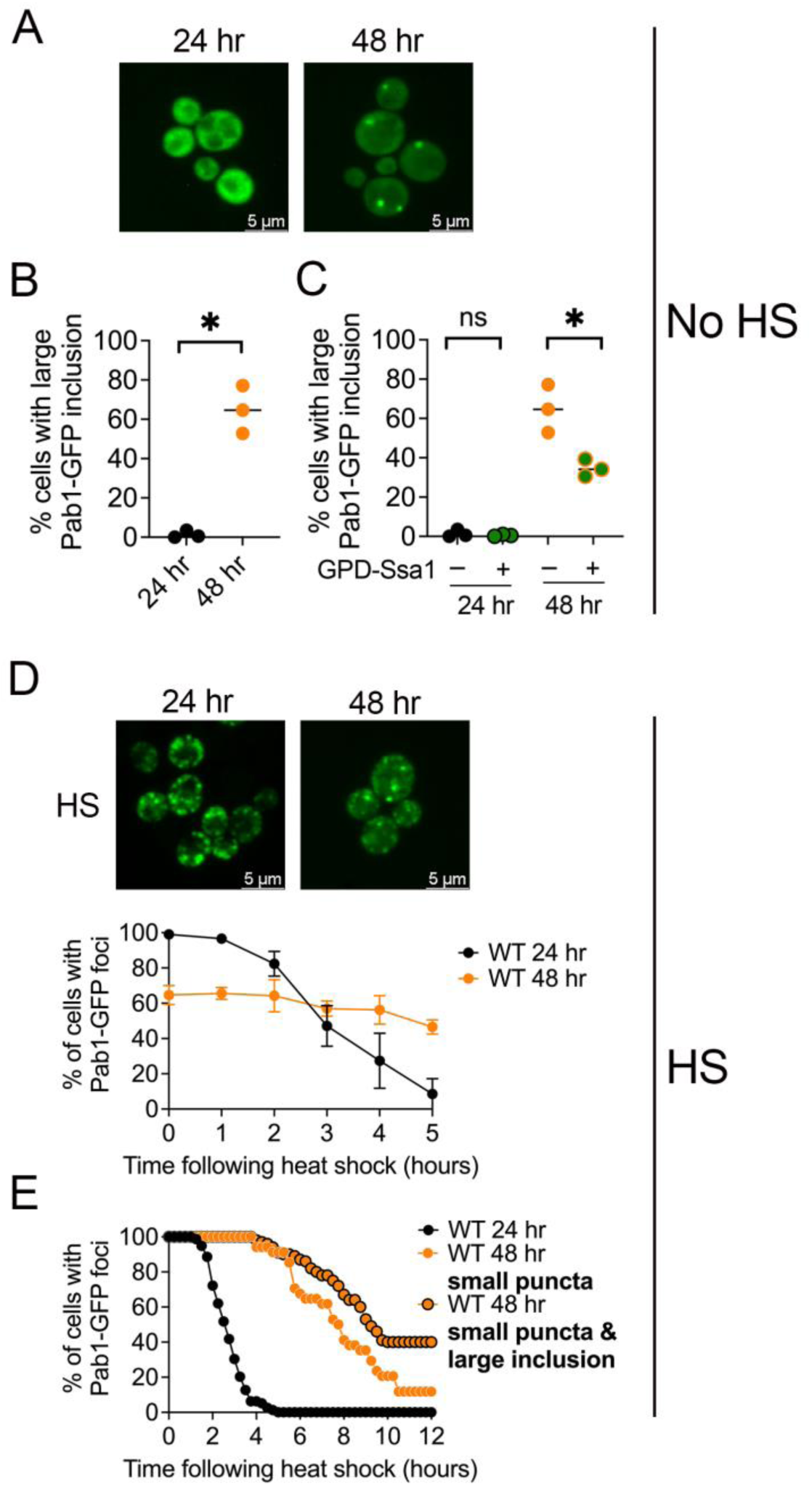
Pab1-GFP forms a large inclusion in saturated cultures which persists following heat shock. A) Pab1-GFP was transformed into a wildtype strain and grown for 24 or 48 hours at 30°C. Widefield images are shown of a paired culture. Other representative images are found in Supplementary figure 6. B) Strains (without heat shock) in A were quantified for the number of cells containing large Pab1 foci. A paired t-test was used to compare changes in the number of cells with Pab1-GFP inclusions (*p<0.01). Horizontal lines are medians. C. *ssa1Δssa2Δ* strains (without heat shock) were transformed with GPD-Ssa1, and cultures were grown for the indicated time. For easier comparison, 24- and 48-hour strains without GPD-Ssa1 (-) are the same as those in B. Unpaired t-test was used in the indicated comparisons (*p<0.02). D) *Top.* Wildtype cultures were heat shocked (42°C for 45 minutes). Other representative images are found in Supplementary figure 6. *Bottom*. After heat shock, widefield images were observed over five hours for the number of cells were scored that lost visible aggregates and exhibited diffuse cytoplasmic fluorescence. Approximately 200 cells from three independent cultures were assessed at the indicated time points after heat shock. All data shown as mean and standard deviation. E) 3D-timelapse microscopy was used to monitor heat shocked 24-hour versus 48-hour cells over twelve hours. 24-hour and 48-hour cells (either containing small puncta or containing both small puncta and large inclusions) were scored for the number of cells that lost any visible aggregates and exhibited diffuse cytoplasmic fluorescence at any given time. The time in which the indicated foci were lost and Pab1-GFP signal became diffuse was quantified and the graph represents the percentage of cells at any given time point that contains Pab1-GFP fluorescent aggregates. Each line indicates anywhere from 37-79 cells monitored by 3D timelapse.

The overexpression of Hsp104 in *ssa1Δssa2Δ* strains does not change the disassembly kinetics of Pab1-GFP, probably due to the already elevated Hsp104 levels in the double mutant (Figure 4A Bottom). Surprisingly, overexpression of GPD-Sis1 exhibited similar disassembly kinetics to the *ssa1Δ* strain or the *ssa1Δssa2Δ* strain overexpressing GPD-Ssa1 (Figure 4A Bottom). Sis1 has been shown to assist Hsp70 in repressing Hsf1 (Feder et al., 2021; Klaips et al., 2020); therefore, it is possible additional Sis1 may increase Hsf1 repression by mediating Hsp70 binding.

Since a portion of *ssa1Δssa2Δ* cells contained the large Pab1-GFP inclusion in addition to the SG puncta, we wanted to determine if these cells exhibited different Pab1-GFP disassembly kinetics. To do this, we discriminated *ssa1Δssa2Δ* cells that either contained 1) *only* normal SG puncta or 2) both the large Pab1 inclusion and SG puncta. To determine the time of Pab1-GFP foci disassembly, we tracked individual cells from the two defined categories over 6 hours using 3D timelapse microscopy following heat shock. All cells from wildtype cultures exhibited normal Pab1-GFP SGs, which showed complete diffuse fluorescence after 4 hours and 30 minutes of recovery (Figure 4B and C). *ssa1Δssa2Δ* cells with only Pab1-GFP SGs resolved faster than wildtype cells, showing all cells with diffuse fluorescence by 3 hours and 45 minutes. This slightly faster recovery could be due to the elevated HSR.

We analyzed the recovery of *ssa1Δssa2Δ* cells that contained both SG and large aggregates. It is important to note that after heat shock, cells containing the large Pab1-GFP inclusions exhibited complete diffuse fluorescence over time (Figure 4B and C). However, the recovery of these cells was slower than wildtype cells or *ssa1Δssa2Δ* cells with only SGs (Figure 4C), showing diffuse fluorescence after six hours. From these studies, we suspect that the elevated HSR in *ssa1Δssa2Δ* cells quickly resolve SGs and provide enough chaperones to slowly dissolve the large Pab1-GFP inclusions after heat shock (Figure 4D), indicating that these large inclusions can be disassembled.

### Ssa1 overexpression reduces the number of cells with large Pab1-GFP in saturated cultures

In *C. elegans* short lifespan mutants, Pab1-GFP forms visible puncta compared to wildtype worms, which have diffuse fluorescence (Midkiff et al., 2022). Since *ssa1Δssa2Δ* mutants also have shorter lifespans (Andersson et al., 2021), the presence of these large inclusions could be a byproduct of aging. We first tested cell viability and Pab1-GFP signal intensity in wildtype cultures that were grown continually over three days in synthetic media. 24-hour cultures were at late log, whereas 48- and 72-hour cultures were at saturation. These saturation conditions mimic cells that are potentially chronologically older. Using methylene blue to assess viability, we found an average of 52.8% of cells were blue following 48 hours of growth, which increased to 85.3% following 72 hours (Figure S5A). Using cells that were expressing Pab1-GFP and stained with the propidium iodide also showed a significant increase in cell death (Figure S5B) that correlated with a decreased Pab1-GFP signal intensity over time (Figure S5C). Due to most of the cells being dead by 72 hours of growth, we decided to only compare 24- and 48-hour cultures. We also assessed whether HSR and chaperone expression is altered in 48-hour cultures. 48-hour cultures exhibited a small but significant increase in HSR compared to 24-hour cultures, which correlated with an increase in endogenously tagged Hsp104-GFP and Sis1-GFP intensity (Figure S5D and E), similar to *ssa1Δssa2Δ* cells (Figure 3).

We next monitored Pab1-GFP aggregation in the two different cultures. Pab1-GFP exhibits cytoplasmic diffuse fluorescence at 24 hours (late log) in wildtype cells under no heat shock (Figure 5A, Figure S6). However, if cultures were allowed to grow continuously for 48 hours, on average of 68.4% of the cells contained a large Pab1-GFP inclusion (Figure 5A and B). If the formation of the Pab1-GFP inclusions in 48-hour cultures was solely due to declining levels of Hsp70s, then the overexpression of Ssa1 should eliminate these inclusions. We found that overexpression of GPD-Ssa1 significantly reduced the Pab1-GFP inclusions to 34.6 or approximately half the amount of cells (Figure 5C), however, it is important to note that the introduction of GPD-Ssa1 did not completely eliminate the presence of these inclusions. These data suggest that presence of Pab1-GFP inclusions in cells from saturated cultures could be due, in part, because of limiting levels of Hsp70.

Next, we asked whether heat shock induced SGs also form in these cultures. While 24-hour cultures show almost all cells contain Pab1-GFP SG puncta upon heat shock, 48-hour cultures show SGs and large inclusions in 64.6% of cells (Figure 5D, timepoint zero; Figure S6). We suspect that the reduce number of cells with Pab1-GFP foci, either large inclusions or SG, is because of the high level of cell death observed in the methylene blue experiments (Figure S5A). Furthermore, since large inclusions are present without heat shock (Figure 5B), we suspect that large Pab1 inclusions in heat shocked cells have formed independently of the SGs. Aggregate disassembly was also different in these two strains. As expected, wildtype 24-hour cultures quickly resolved Pab1-GFP foci, with only 8.5% of cells containing aggregates after five hours. Conversely, heat shocked 48-hour cultures appeared to maintain approximately 60-70% of cells with visible Pab1-GFP foci over time and do not recover by five hours (Figure 5D).

Since it was possible that our timeframes of tracking disassembly were not long enough, we performed 3D-timelapse microscopy to track individual cells over twelve hours after heat shock. Similar to our *ssa1Δssa2Δ* recovery studies in Figure 4C, we categorized cells as 1) 24-hour cells with SGs, 2) 48-hour cultures that contain *only* SGs, or 3) 48-hour cultures that contain both SGs and large Pab1 inclusions. Because cells lacking foci were not followed during the timelapse, each of the above categories was normalized to 100% to more easily assay their relative rate of recovery. In all cell types, we monitored the time in which cells transitioned from containing any Pab1-GFP foci to diffuse fluorescence. All cells from 24-hour cultures exhibited disassembled Pab1-GFP SGs by 5 hours (Figure 5E). Cells from 48-hour cultures exhibited a delay in disassembly, regardless of which category of Pab1-GFP foci they exhibited (Figure 5E). In 48-hour cultures, 11.8% cells contained small SG puncta after 12 hours. Conversely, 40% of cells that contained both small SG puncta and large inclusions did not recover after 12 hours. (Figure 5E). While both saturated cells (Figure 5E) and *ssa1Δssa2Δ* cells containing a Pab1-GFP inclusion (Figure 4C) exhibited slower foci recovery, the kinetics were on dramatically different timescales. We suspect these age-associated inclusions are not limited by Hsp70 levels alone and could be different from those observed in *ssa1Δssa2Δ* cells.

## DISCUSSION

### Hsp70 and limiting unwanted protein aggregation

Previous studies with model aggregation proteins, such as NES-LuciTs, or mutant yeast Ubc9Ts, *guk1-7*, and *gus1-3* proteins, show that these proteins form inclusions in *ssa1Δssa2Δ* <u>cells</u> (Andersson et al., 2021; Rolli et al., 2024; Shiber et al., 2013). Since these are either heterologous or mutant proteins, the role of Hsp70 in limiting aggregation of endogenous proteins was poorly explored. Our studies, using the endogenous Pab1 protein that is intrinsically disordered and is associated with stress granules upon transient heat shock, show that this protein forms large inclusions in *ssa1Δssa2Δ* cells under normal conditions. These inclusions were absent in *ssa1Δssa2Δ* cells overexpressing either Ssa1, Sis1, or Hsp104 under no heat shock conditions, suggesting that increasing amounts of any of these chaperones can limit Pab1 from forming inclusions under normal conditions, and Sis1 and Hsp104 are possibly able to compensate for the lack of Ssa1 and Ssa2.

Interestingly during hat shock, large Pab1-GFP inclusions formed in *ssa1Δssa2Δ* strains despite the overexpression of Hsp104 or Sis1. However, excess of Ssa1 appeared to limit these large inclusions during heat shock (Figure 1), suggesting that Ssa1 plays a prominent role in limiting unwanted aggregation of proteins that naturally aggregate in response to stress, such as Pab1. Our data suggests these Hsp70s may have general roles in limiting unwanted protein aggregation, and we suspect that the IDR region of Pab1 may be more prone to misfolding and aggregation when insufficient levels of Hsp70s are present.

### Elevated HSR in ssa1Δssa2Δ

Hsp70s play numerous roles in protein refolding, and disassembly (Kampinga and Craig, 2010; Lotz et al., 2019; Ruger-Herreros et al., 2024). In addition, Hsp70s also play a role in inactivating Hsf1, thereby keeping the HSR low (Krakowiak et al., 2018). However, upon heat shock, changes in proteostasis titrate Hsp70 away from Hsf1, allowing the trimerization of Hsf1 and subsequent activation of the heat shock genes (Masser et al., 2019). Meanwhile during stress recovery, chaperones such as Hsp104 mediate the disassembly of SGs during recovery from heat shock. In our studies, SG recovery in wildtype cell populations happen over five hours. Interestingly, the recovery kinetics of SGs is faster in *ssa1Δssa2Δ* strains (Figure 4A). We suspect that under normal conditions, *ssa1Δssa2Δ* cells are in a constant state of elevated stress because the low level of Hsp70s in *ssa1Δssa2Δ* are insufficient to repress Hsf1 activity and thereby cause the HSR to remain “on”. Therefore, constitutive activation of Hsf1 and the continuous upregulation other chaperones like Hsp104 and Sis1 (Figure 3), could prime *ssa1Δssa2Δ* cells for fast Pab1-GFP SG disassembly after heat stress (Figure 4A). However, there could be a cost for this constitutive HSR, resulting in shorter lifespan and issues with cell growth (Figure S4). Interestingly, our results showed that loss of Ssa1 or Ssa2 alone (*ssa1Δ* vs. *ssa1Δssa2Δ* expressing GPD-Ssa1) results in slower recovery (Figure 4A). These strains could have sufficient amounts of Hsp70 to repress Hsf1 under normal conditions, yet insufficient amounts of Hsp70 to ensure chaperone-mediated disassembly under heat stress. In fact, the similar HSR levels in *ssa1Δ* and wildtype strains (Figure 3A), could potentially explain these observations.

Although quick recovery from stress is thought to be advantageous, we did not see a correlation between the fast recovery of small Pab1-GFP foci after heat shock and cell growth in *ssa1Δssa2Δ* strains (Figure S4). In fact, despite the fast recovery, cells appeared to be sicker than wildtype. This phenomenon has also been observed for other chaperones. For example, chronic activation of HSR by Sis1 overexpression leads to cell hypersensitivity to stress (Klaips et al., 2020; Lamech and Haynes, 2015; Roth et al., 2014). By parsing out *ssa1Δssa2Δ* cells containing a large Pab1-GFP inclusion or only normal Pab1-GFP SG puncta, we find cells with large inclusions take longer to disassemble following heat shock (Figure 4C). Other studies have proposed large inclusions formed by heterologous or temperature sensitive mutant proteins when Hsp70 is lost are more solid and less dynamic (Andersson et al., 2021; Grosfeld et al., 2023; Rolli et al., 2024; Shiber et al., 2013). Since Pab1 is a core SG protein, the protein may form more stable interactions with other proteins found within the large Pab1-GFP inclusions in *ssa1Δssa2Δ* strains. These stable inclusions may be resistant to chaperone-mediated clearance. However, the increase in chaperones levels after heat shock may be sufficient for the eventual, albeit slower, disassembly of these inclusions.

### Link between Pab1-GFP inclusions and Hsp70 during aging

*ssa1Δssa2Δ* cells have a shortened lifespan (Andersson et al., 2021). Since Pab1-GFP forms inclusions under normal conditions in *ssa1Δssa2Δ,* we investigated whether Pab1-GFP would form inclusions in wildtype cultures that have been grown to saturation. While cell division and metabolism are robust in log phase cultures, saturated cultures usually have increased chronological aging, have low nutrient availability and metabolism as well as lower ATP levels (Bilinski et al., 2017; Gray et al., 2004). In addition, acidification of the media is also observed in saturated cultures. Similar to *ssa1Δssa2Δ* cells, Pab1-GFP formed large inclusions in saturated wildtype cultures without heat shock (Figure 5). Overexpression of Ssa1 reduced, but did not eliminate, the number of cells that contained these large inclusions. We suspect aging cells are facing proteostasis collapse, resulting in insufficient levels of Hsp70 available to prevent unwanted aggregation of proteins, including those which have intrinsically disordered regions. Alternatively, Pab1 may be more prone to aggregation as the cell conditions change over time. Since Pab1 demixes in acidic conditions (Riback et al., 2017), it is also possible that the change in pH in saturated cultures could be sufficient to drive Pab1, and possibly other IDR containing proteins, into inclusions. However, introduction of extra Ssa1 appears to lessen the amount of Pab1-GFP inclusions in these saturated cultures (Figure 5B). indicating that Hsp70s play an important role in regulating Pab1 aggregation.

While we observed a Hsp70 dependence in the formation of large Pab1-GFP inclusions, disassembly told a different story. Cells containing only SGs appeared to recover faster in *ssa1Δssa2Δ* compared to wildtype (Figure 4C), whereas saturated cultures showed a severe recovery delay in cells containing only SGs compared to 24-hour cultures (Figure 5E). Since Hsp104 plays a central role in SG disassembly (Kroschwald et al., 2015), we suspect that the elevated HSR and increased levels of Hsp104 in *ssa1Δssa2Δ* contributes to faster SG disassembly, however, in saturated cultures, the accumulation of misfolded proteins may overwhelm the chaperone machinery. The chaperone availability of Ssa1, as well as other chaperones like Ydj1, has been shown to be reduced in aging yeast cells, despite an increase in Hsp104 levels (Moreno et al., 2019). We suspect that the slower recovery is not necessarily linked to Hsp70 alone, but availability of multiple chaperones that are required for the disassembly process.

## MATERIALS AND METHODS

### Strains, plasmids, and cultivation procedures

All strains (Supplementary Table 1) were in the 74D-694 (*Mat**a** ade1-14 leu2-3,112 his3-Δ200 trp1-289 ura3-52* [*pin*^-^] [*psi*^-^]) background (Chernoff et al., 1995), and grown at 30°C using standard synthetic media and cultivation procedures (Sherman et al., 1986). Note that all strains in this study were [*pin^-^*] and therefore should not contain or induce prion aggregates (Derkatch et al., 1997; Zhou et al., 2001). For no heat shock and heat shock studies, a single culture was split after growth at 30°C. One sample was incubated at room temperature (approximately 22-25°C) and was considered our no or without heat shock condition. The other sample was incubated at 42°C for 45 minutes. Note that 42°C degrees was chosen since this genetic background had viability issues beyond 44°C. Strains transformed with plasmids (Supplemental Table 2) were maintained on synthetic complete media lacking the specific amino acid(s) to maintain the plasmid. *ssa1* deletion (*ssa1*Δ) strain in the 74D-694 background was disrupted with a HIS3 cassette. *SSA2* was disrupted in the *ssa1Δ* strain by introduction of a *NatR* cassette.

Pab1-GFP (3284, 3285) was a gift from Roy Parker (Brengues and Parker, 2007). The Pab1-mCherry plasmid was constructed as follows: primers (AM583 5’-TACATAGAGCGTCGACCAGCTTGCTCAGTTTGTTGTTCTTGC-3’ & AM586 5’-GACCATATAAGAATTCATACTGTATGAAGCC-3’) were used to amplify Pab1 from BY4741 genomic DNA, followed by traditional restriction enzyme-mediated cloning using AccI and EcoRI into pGEM-T Easy that already contained mCherry with a short N-terminal linker peptide. The Pab1-mCherry fusion construct was sequence verified, and then subcloned into pRS314 plasmid to provide Trp1 selection. Ssa1 driven by a *GPD* promoter (p3302), and Sis1 driven by a *GPD* promoter (p3281) were constructed by amplifying the open reading frames from genomic DNA, and using the Gateway cloning system and yeast destination vectors (Alberti et al., 2007). Hsp104 driven by its native promoter (p3110) (Jackrel et al., 2014), or Sup35 PrD-GFP (which is the N-terminal 1-254 amino acids of Sup35 fused to GFP) driven by a copper-inducible (*CUP1)* promoter (Zhou et al., 2001) were used as indicated. To integrate the HSE-YFP reporter (p3357) (Zheng et al., 2016) into wildtype, *ssa1Δ*, and *ssa1Δ ssa2Δ* strains, p3357 was linearized with PmeI, transformed into indicated strains, and selected on media lacking leucine to generate M703, M771, and M707, respectively. To integrate nuclear marker Nab2-mCherry (p3272) into wildtype and *ssa1Δ ssa2Δ* strains, p3272 was linearized with AvrII, transformed into indicated strains, and selected on media lacking tryptophan to generate M761 and M757, respectively (Zhu et al., 2019). To integrate vacuolar marker Vph1-mCherry (p3273) into wildtype and *ssa1Δssa2Δ* strains, 3273 was linearized with PmlI, transformed into indicated strains, and selected on media lacking tryptophan to generate M762 and M763, respectively (Zhu et al., 2019).

For serial dilution plates, all strains were grown in liquid culture overnight to late log and plated at five-fold serial dilutions on indicated selective media. Plates were incubated at 30°C for 2-3 days. For wildtype strains grown over 72 hours, strains were inoculated in 5-ml of plasmid selective synthetic media and grown at 30°C with shaking for the indicated times. 24-hour cultures had an OD_600_ of approximately 0.9-1.2, which is considered late log. 48- and 72-hour cultures had OD_600_ readings over 1.4 in synthetic media.

### Widefield fluorescent microscopy

To characterize Pab1-GFP aggregate morphology and disassembly, wildtype, *ssa1Δ*, and *ssa1Δ ssa2Δ* strains containing a Pab1-GFP plasmid (p3284 or p3285) alone or with an additional chaperone overexpression plasmid (p3302, p3110, p3281) were grown overnight to late log phase at 30°C. Cultures were either heat shocked for 45 minutes at 42°C or left at room temperature for the same period of time, and 3D images of fields were acquired with Leica DMI 6000 fluorescent deconvolution microscope (63X, 1.4 NA) and captured with a Leica K5 camera. Similarly for disassembly, indicated cultures were heat shocked for 45 minutes at 42°C and 3D field images were acquired every hour following heat shock for five hours. Images were captured as above, and the number of cells with Pab1-GFP granules were quantified using LASX software.

For 3D timelapse microscopy to follow individual cells over time, indicated strains were grown overnight to late log at 30°C. Cells were then heat shocked for 45 minutes at 42°C and 200 µL of culture was immediately added to a concanavalin A-coated Ibidi 8-well glass bottom slide. Cells were incubated with 300 µL fresh selective media, and 3D images were captured every 15 minutes for 3 hours, using a 100 X, NA 1.44 objective. Images were processed with LASX software. All images are shown with 3D deconvolution (Media Cybernetics) and maximum projection.

Methylene blue was used to stain dead cells. 1.5 µL of resuspended cells were placed onto glass microscope slides, and 1.5 μL of 1X methylene blue were added directly to the sample. Cell mixtures were allowed to incubate for 5 minutes before 3D brightfield images were acquired with Leica DMI 6000 fluorescent deconvolution microscope (63X, 1.4 NA) and captured with a Leica K5 camera.

### Spinning disk confocal microscopy

For higher resolution, spinning disk confocal microscopy was used. Cultures were either left untreated or heat shocked for 45 minutes at 42°C. Images were acquired with a Zeiss AxioObserver, CrestOptics Cicero spinning-disk confocal system (100X, 1.44 NA), and captured with ORCA Flash CMOS camera. Images were processed using VisiView Software.

### Flow cytometry

Wildtype, *ssa1Δ*, and *ssa1Δ ssa2Δ* strains integrated with the HSE-YFP reporter were grown overnight to late log phase at 30°C. Cultures were either left untreated or heat shocked for 45 minutes at 42°C. Flow cytometry was performed using a Cytoflex Flow Cytometer (Beckman Coulter) to measure HSE-YFP intensity (FITC filter). 100,000 cells were counted per sample. Histograms were generated using CytExpert Software. Aging experiments were conducted similarly except samples were treated with propidium iodide (PI) to stain and gate out dead cells. 400 µL of culture was centrifuged at 3000 rpm for 3 minutes. The pellet was resuspended in 400 µL PBS and 4 µg/ml of PI. Cells were incubated on ice in the dark for 15 minutes. 100,000 cells were counted per sample. Viable cells were gated from dead cells based upon low ECD signal, leaving only live cells reported within the FITC-A peaks.

### Western blotting

Wildtype and *ssa1Δ ssa2Δ* strains containing a Pab1-GFP plasmid (3284) were grown overnight to late log phase at 30°C. Cultures were either left untreated or heat shocked for 45 minutes at 42°C, and cell lysates were prepared by glass bead cold lysis with 1x lysis buffer (Knier et al., 2022; Sharma et al., 2017), protease inhibitor cocktail (Sigma) and PMSF (Sigma) in the Cryolys Evolution (Bertin Technologies). Cells were lysed by shaking at 6300 RPM every 30 seconds while being cooled to 1°C. Debris was cleared via centrifugation (1000 rpm). 100 µg of cell extracts was treated with 2% SDS sample buffer (Knier et al., 2022) and boiled prior to resolving on a 10% acrylamide gel to analyze protein steady state levels. Blots were subjected to normal western blotting procedures. Antibody conditions used are described in supplemental table 3. Antibody signal was captured with the iBright 1500 (ThermoFisher) and quantified in ImageJ. All signals were normalized to tubulin loading control.

## ABBREVIATIONS

(HSR): Heat shock response
(SG): stress granule
(IDR): intrinsically disordered region
(JDP): J-Domain proteins

## Acknowledgements

The authors would like to thank David Pincus (UChicago), Roy Parker (UC Boulder) and James Shorter (UPenn) for plasmids, and Elizabeth Craig (UW-Madison) for antibodies used in these studies. Strains and plasmids were a kind gift from Susan Liebman, and the BE4 (Sup35C) antibody was a gift from Viravan Prapapanich and Susan Liebman. This work was supported by the National Science Foundation (MCB 2127616) and the National Institutes of Health (GM155860) to ALM. HEB was supported by the Marquette Raynor Fellowship, ASK was supported by the Schmitt Leadership Fellowship, and CMR was supported by the GAANN fellowship.

## Notes

### Competing Interest Statement

The authors have declared no competing interest.

## REFERENCES

Alberti, S., A.D. Gitler, and S. Lindquist. 2007. A suite of Gateway cloning vectors for high-throughput genetic analysis in Saccharomyces cerevisiae. Yeast. 24:913–919. PMC2190539

Anckar, J., and L. Sistonen. 2011. Regulation of HSF1 function in the heat stress response: implications in aging and disease. Annu Rev Biochem. 80:1089–1115. PMID21417720

Anderson, P., and N. Kedersha. 2006. RNA granules. J Cell Biol. 172:803–808. PMC2063724

Andersson, R., A.M. Eisele-Burger, S. Hanzen, K. Vielfort, D. Oling, F. Eisele, G. Johansson, T. Gustafsson, K. Kvint, and T. Nystrom. 2021. Differential role of cytosolic Hsp70s in longevity assurance and protein quality control. PLoS genetics. 17:e1008951. PMC7822560

Bilinski, T., A. Bylak, and R. Zadrag-Tecza. 2017. The budding yeast Saccharomyces cerevisiae as a model organism: possible implications for gerontological studies. Biogerontology. 18:631–640. PMC5514200

Boorstein, W.R., and E.A. Craig. 1990. Structure and regulation of the SSA4 HSP70 gene of Saccharomyces cerevisiae. J Biol Chem. 265:18912–18921. PMID2121731

Brambilla, M., F. Martani, S. Bertacchi, I. Vitangeli, and P. Branduardi. 2019. The Saccharomyces cerevisiae poly (A) binding protein (Pab1): Master regulator of mRNA metabolism and cell physiology. Yeast. 36:23–34. PMID30006991

Brengues, M., and R. Parker. 2007. Accumulation of polyadenylated mRNA, Pab1p, eIF4E, and eIF4G with P-bodies in Saccharomyces cerevisiae. Mol Biol Cell. 18:2592–2602. PMC1924816

Buchan, J.R., and R. Parker. 2009. Eukaryotic stress granules: the ins and outs of translation. Mol Cell. 36:932–941. PMC2813218

Buchholz, H.E., J.E. Dorweiler, S. Guereca, B.T. Wisniewski, J. Shorter, and A.L. Manogaran. 2024. The middle domain of Hsp104 can ensure substrates are functional after processing. PLoS genetics. 20:e1011424. PMC11478891

Chakrabortee, S., J.S. Byers, S. Jones, D.M. Garcia, B. Bhullar, A. Chang, R. She, L. Lee, B. Fremin, S. Lindquist, and D.F. Jarosz. 2016. Intrinsically Disordered Proteins Drive Emergence and Inheritance of Biological Traits. Cell. 167:369–381 e312. PMC5066306

Chernoff, Y.O., S.L. Lindquist, B. Ono, S.G. Inge-Vechtomov, and S.W. Liebman. 1995. Role of the chaperone protein Hsp104 in propagation of the yeast prion-like factor [psi+]. Science. 268:880–884. PMID7754373

Chernova, T.A., K.D. Wilkinson, and Y.O. Chernoff. 2017. Prions, Chaperones, and Proteostasis in Yeast. Cold Spring Harb Perspect Biol. 9. PMC5287078

Clerico, E.M., J.M. Tilitsky, W. Meng, and L.M. Gierasch. 2015. How hsp70 molecular machines interact with their substrates to mediate diverse physiological functions. J Mol Biol. 427:1575–1588. PMC4440321

Comyn, S.A., B.P. Young, C.J. Loewen, and T. Mayor. 2016. Prefoldin Promotes Proteasomal Degradation of Cytosolic Proteins with Missense Mutations by Maintaining Substrate Solubility. PLoS Genet. 12:e1006184. PMC4957761

Craig, E.A., and K. Jacobsen. 1984. Mutations of the heat inducible 70 kilodalton genes of yeast confer temperature sensitive growth. Cell. 38:841–849. PMID6386178

Derkatch, I.L., M.E. Bradley, P. Zhou, Y.O. Chernoff, and S.W. Liebman. 1997. Genetic and environmental factors affecting the de novo appearance of the [PSI+] prion in Saccharomyces cerevisiae. Genetics. 147:507–519. PMC1208174

Escusa-Toret, S., W.I. Vonk, and J. Frydman. 2013. Spatial sequestration of misfolded proteins by a dynamic chaperone pathway enhances cellular fitness during stress. Nat Cell Biol. 15:1231–1243. PMC4121856

Feder, Z.A., A. Ali, A. Singh, J. Krakowiak, X. Zheng, V.P. Bindokas, D. Wolfgeher, S.J. Kron, and D. Pincus. 2021. Subcellular localization of the J-protein Sis1 regulates the heat shock response. J Cell Biol. 220. PMC7748816

Franzmann, T.M., M. Jahnel, A. Pozniakovsky, J. Mahamid, A.S. Holehouse, E. Nuske, D. Richter, W. Baumeister, S.W. Grill, R.V. Pappu, A.A. Hyman, and S. Alberti. 2018. Phase separation of a yeast prion protein promotes cellular fitness. Science. 359. PMID29301985

Gidalevitz, T., V. Prahlad, and R.I. Morimoto. 2011. The stress of protein misfolding: from single cells to multicellular organisms. Cold Spring Harb Perspect Biol. 3. PMC3098679

Glover, J.R., and S. Lindquist. 1998. Hsp104, Hsp70, and Hsp40: a novel chaperone system that rescues previously aggregated proteins. Cell. 94:73–82. PMID9674429

Gray, J.V., G.A. Petsko, G.C. Johnston, D. Ringe, R.A. Singer, and M. Werner-Washburne. 2004. “Sleeping beauty”: quiescence in Saccharomyces cerevisiae. Microbiol Mol Biol Rev. 68:187–206. PMC419917

Grimes, B., W. Jacob, A.R. Liberman, N. Kim, X. Zhao, D.C. Masison, and L.E. Greene. 2023. The Properties and Domain Requirements for Phase Separation of the Sup35 Prion Protein In Vivo. Biomolecules. 13. PMC10526957

Grizel, A.V., N.A. Gorsheneva, J.B. Stevenson, J. Pflaum, F. Wilfling, A.A. Rubel, and Y.O. Chernoff. 2024. Osmotic stress induces formation of both liquid condensates and amyloids by a yeast prion domain. J Biol Chem. 300:107766. PMID39276934

Grosfeld, E.V., A.Y. Beizer, A.A. Dergalev, M.O. Agaphonov, and A.I. Alexandrov. 2023. Fusion of Hsp70 to GFP Impairs Its Function and Causes Formation of Misfolded Protein Deposits under Mild Stress in Yeast. Int J Mol Sci. 24. PMC10454418

Gross, D.S., C.C. Adams, K.E. English, K.W. Collins, and S. Lee. 1990. Promoter function and in situ protein/DNA interactions upstream of the yeast HSP90 heat shock genes. Antonie Van Leeuwenhoek. 58:175–186. PMID2256678

Hasin, N., S.A. Cusack, S.S. Ali, D.A. Fitzpatrick, and G.W. Jones. 2014. Global transcript and phenotypic analysis of yeast cells expressing Ssa1, Ssa2, Ssa3 or Ssa4 as sole source of cytosolic Hsp70-Ssa chaperone activity. BMC Genomics. 15:194. PMC4022180

Horst, M., E.C. Knecht, and P.V. Schu. 1999. Import into and degradation of cytosolic proteins by isolated yeast vacuoles. Mol Biol Cell. 10:2879–2889. PMC25526

Howard, M.K., B.S. Sohn, J. von Borcke, A. Xu, and M.E. Jackrel. 2020. Functional analysis of proposed substrate-binding residues of Hsp104. PLoS One. 15:e0230198. PMC7064214

Hoyle, N.P., L.M. Castelli, S.G. Campbell, L.E. Holmes, and M.P. Ashe. 2007. Stress-dependent relocalization of translationally primed mRNPs to cytoplasmic granules that are kinetically and spatially distinct from P-bodies. J Cell Biol. 179:65–74. PMC2064737

Jackrel, M.E., M.E. DeSantis, B.A. Martinez, L.M. Castellano, R.M. Stewart, K.A. Caldwell, G.A. Caldwell, and J. Shorter. 2014. Potentiated Hsp104 variants antagonize diverse proteotoxic misfolding events. Cell. 156:170–182. PMC3909490

Jackrel, M.E., and J. Shorter. 2014. Potentiated Hsp104 variants suppress toxicity of diverse neurodegenerative disease-linked proteins. Dis Model Mech. 7:1175–1184. PMC4174528

Jain, S., J.R. Wheeler, R.W. Walters, A. Agrawal, A. Barsic, and R. Parker. 2016. ATPase-Modulated Stress Granules Contain a Diverse Proteome and Substructure. Cell. 164:487–498. PMC4733397

Jawed, A., C.T. Ho, T. Grousl, A. Shrivastava, T. Ruppert, B. Bukau, and A. Mogk. 2022. Balanced activities of Hsp70 and the ubiquitin proteasome system underlie cellular protein homeostasis. Front Mol Biosci. 9:1106477. PMC9845930

Jones, L., K. Tedrick, A. Baier, M.R. Logan, and G. Eitzen. 2010. Cdc42p is activated during vacuole membrane fusion in a sterol-dependent subreaction of priming. J Biol Chem. 285:4298–4306. PMC2836034

Kaganovich, D., R. Kopito, and J. Frydman. 2008. Misfolded proteins partition between two distinct quality control compartments. Nature. 454:1088–1095. PMID18756251

Kampinga, H.H., and E.A. Craig. 2010. The HSP70 chaperone machinery: J proteins as drivers of functional specificity. Nat Rev Mol Cell Biol. 11:579–592. PMC3003299

Kedersha, N., and P. Anderson. 2002. Stress granules: sites of mRNA triage that regulate mRNA stability and translatability. Biochem Soc Trans. 30:963–969. PMID12440955

Klaips, C.L., M.H.M. Gropp, M.S. Hipp, and F.U. Hartl. 2020. Sis1 potentiates the stress response to protein aggregation and elevated temperature. Nat Commun. 11:6271. PMC7722728

Knier, A.S., E.E. Davis, H.E. Buchholz, J.E. Dorweiler, L.E. Flannagan, and A.L. Manogaran. 2022. The yeast molecular chaperone, Hsp104, influences Transthyretin (TTR) aggregate formation. Front. Mol. Neurosci. doi: 10.3389/fnmol.2022.1050472. PMC9802906

Krakowiak, J., X. Zheng, N. Patel, Z.A. Feder, J. Anandhakumar, K. Valerius, D.S. Gross, A.S. Khalil, and D. Pincus. 2018. Hsf1 and Hsp70 constitute a two-component feedback loop that regulates the yeast heat shock response. eLife. 7. PMC5809143

Kroschwald, S., S. Maharana, D. Mateju, L. Malinovska, E. Nuske, I. Poser, D. Richter, and S. Alberti. 2015. Promiscuous interactions and protein disaggregases determine the material state of stress-inducible RNP granules. eLife. 4:e06807. PMC4522596

Lamech, L.T., and C.M. Haynes. 2015. The unpredictability of prolonged activation of stress response pathways. J Cell Biol. 209:781–787. PMC4477854

Lindquist, S. 1986. The heat-shock response. Annual review of biochemistry. 55:1151–1191. PMID2427013

Lotz, S.K., L.E. Knighton, Nitika, G.W. Jones, and A.W. Truman. 2019. Not quite the SSAme: unique roles for the yeast cytosolic Hsp70s. Curr Genet. 65:1127–1134. PMC7262668

Masser, A.E., W. Kang, J. Roy, J. Mohanakrishnan Kaimal, J. Quintana-Cordero, M.R. Friedlander, and C. Andreasson. 2019. Cytoplasmic protein misfolding titrates Hsp70 to activate nuclear Hsf1. eLife. 8. PMC6779467

Midkiff, D.F., J. Huayta, J.D. Lichty, J.P. Crapster, and A. San-Miguel. 2022. Identifying C. elegans lifespan mutants by screening for early-onset protein aggregation. iScience. 25:105460. PMC9664360

Moreno, D.F., K. Jenkins, S. Morlot, G. Charvin, A. Csikasz-Nagy, and M. Aldea. 2019. Proteostasis collapse, a hallmark of aging, hinders the chaperone-Start network and arrests cells in G1. eLife. 8. PMC6744273

Oling, D., F. Eisele, K. Kvint, and T. Nystrom. 2014. Opposing roles of Ubp3-dependent deubiquitination regulate replicative life span and heat resistance. EMBO J. 33:747–761. PMC4000091

Pessa, J.C., J. Joutsen, and L. Sistonen. 2024. Transcriptional reprogramming at the intersection of the heat shock response and proteostasis. Mol Cell. 84:80–93. PMID38103561

Riback, J.A., C.D. Katanski, J.L. Kear-Scott, E.V. Pilipenko, A.E. Rojek, T.R. Sosnick, and D.A. Drummond. 2017. Stress-Triggered Phase Separation Is an Adaptive, Evolutionarily Tuned Response. Cell. 168:1028–1040 e1019. PMC5401687

Rolli, S., C.A. Langridge, and E.M. Sontag. 2024. Clearing the JUNQ: the molecular machinery for sequestration, localization, and degradation of the JUNQ compartment. Front Mol Biosci. 11:1427542. PMC11372896

Roth, D.M., D.M. Hutt, J. Tong, M. Bouchecareilh, N. Wang, T. Seeley, J.F. Dekkers, J.M. Beekman, D. Garza, L. Drew, E. Masliah, R.I. Morimoto, and W.E. Balch. 2014. Modulation of the maladaptive stress response to manage diseases of protein folding. PLoS Biol. 12:e1001998. PMC4236052

Ruger-Herreros, C., L. Svoboda, A. Mogk, and B. Bukau. 2024. Role of J-domain Proteins in Yeast Physiology and Protein Quality Control. J Mol Biol. 436:168484. PMID38331212

Sharma, J., B.T. Wisniewski, E. Paulson, J.O. Obaoye, S.J. Merrill, and A.L. Manogaran. 2017. De novo [PSI +] prion formation involves multiple pathways to form infectious oligomers. Sci Rep. 7:76. PMC5427932

Shattuck, J.E., K.R. Paul, S.M. Cascarina, and E.D. Ross. 2019. The prion-like protein kinase Sky1 is required for efficient stress granule disassembly. Nat Commun. 10:3614. PMC6688984

Sherman, F., G.R. Fink, and J.B. Hicks. 1986. *Methods in Yeast Genetics*. Cold Spring Harbor Lab., Plainview, New York.

Shiber, A., W. Breuer, M. Brandeis, and T. Ravid. 2013. Ubiquitin conjugation triggers misfolded protein sequestration into quality control foci when Hsp70 chaperone levels are limiting. Mol Biol Cell. 24:2076–2087. PMC3694792

Shorter, J., and S. Lindquist. 2008. Hsp104, Hsp70 and Hsp40 interplay regulates formation, growth and elimination of Sup35 prions. EMBO J. 27:2712–2724. PMC2572177

Sorger, P.K., and H.C. Nelson. 1989. Trimerization of a yeast transcriptional activator via a coiled-coil motif. Cell. 59:807–813. PMID2686840

Sweeny, E.A., M.E. Jackrel, M.S. Go, M.A. Sochor, B.M. Razzo, M.E. DeSantis, K. Gupta, and J. Shorter. 2015. The Hsp104 N-terminal domain enables disaggregase plasticity and potentiation. Mol Cell. 57:836–849. PMC4623595

Swisher, K.D., and R. Parker. 2010. Localization to, and effects of Pbp1, Pbp4, Lsm12, Dhh1, and Pab1 on stress granules in Saccharomyces cerevisiae. PLoS One. 5:e10006. PMC2848848

Verghese, J., J. Abrams, Y. Wang, and K.A. Morano. 2012. Biology of the heat shock response and protein chaperones: budding yeast (Saccharomyces cerevisiae) as a model system. Microbiol Mol Biol Rev. 76:115–158. PMC3372250

Voellmy, R., and F. Boellmann. 2007. Chaperone regulation of the heat shock protein response. Adv Exp Med Biol. 594:89–99. PMID17205678

Wallace, E.W., J.L. Kear-Scott, E.V. Pilipenko, M.H. Schwartz, P.R. Laskowski, A.E. Rojek, C.D. Katanski, J.A. Riback, M.F. Dion, A.M. Franks, E.M. Airoldi, T. Pan, B.A. Budnik, and D.A. Drummond. 2015. Reversible, Specific, Active Aggregates of Endogenous Proteins Assemble upon Heat Stress. Cell. 162:1286–1298. PMC4567705

Walters, R.W., D. Muhlrad, J. Garcia, and R. Parker. 2015. Differential effects of Ydj1 and Sis1 on Hsp70-mediated clearance of stress granules in Saccharomyces cerevisiae. RNA. 21:1660–1671. PMC4536325

Werner-Washburne, M., D.E. Stone, and E.A. Craig. 1987. Complex interactions among members of an essential subfamily of hsp70 genes in Saccharomyces cerevisiae. Mol Cell Biol. 7:2568–2577. PMC365392

Wheeler, J.R., T. Matheny, S. Jain, R. Abrisch, and R. Parker. 2016. Distinct stages in stress granule assembly and disassembly. Elife. 5. PMC5014549

Winkler, J., J. Tyedmers, B. Bukau, and A. Mogk. 2012. Hsp70 targets Hsp100 chaperones to substrates for protein disaggregation and prion fragmentation. J Cell Biol. 198:387–404. PMC3413357

Yoo, H., J.A.M. Bard, E.V. Pilipenko, and D.A. Drummond. 2022. Chaperones directly and efficiently disperse stress-triggered biomolecular condensates. Mol Cell. 82:741–755 e711. PMC8857057

Young, M.R., and E.A. Craig. 1993. Saccharomyces cerevisiae HSP70 heat shock elements are functionally distinct. Mol Cell Biol. 13:5637–5646. PMC360292

Zheng, X., J. Krakowiak, N. Patel, A. Beyzavi, J. Ezike, A.S. Khalil, and D. Pincus. 2016. Dynamic control of Hsf1 during heat shock by a chaperone switch and phosphorylation. eLife. 5. PMC5127643

Zhou, P., I.L. Derkatch, and S.W. Liebman. 2001. The relationship between visible intracellular aggregates that appear after overexpression of Sup35 and the yeast prion-like elements [PSI(+)] and [PIN(+)]. Mol Microbiol. 39:37–46. PMID11123686

Zhu, J., Z.T. Zhang, S.W. Tang, B.S. Zhao, H. Li, J.Z. Song, D. Li, and Z. Xie. 2019. A Validated Set of Fluorescent-Protein-Based Markers for Major Organelles in Yeast (Saccharomyces cerevisiae). mBio. 10. PMC6722415

