## Supplemental figures and Tables for "Hsp70 chaperones, Ssa1 and Ssa2, limit poly(A) binding protein aggregation"

**RUNNING HEAD** Hsp70 limits Pab1 aggregation

**ABBREVIATIONS** Heat shock response (HSR), stress granule (SG), intrinsically disordered region (IDR), J-Domain proteins (JDP)

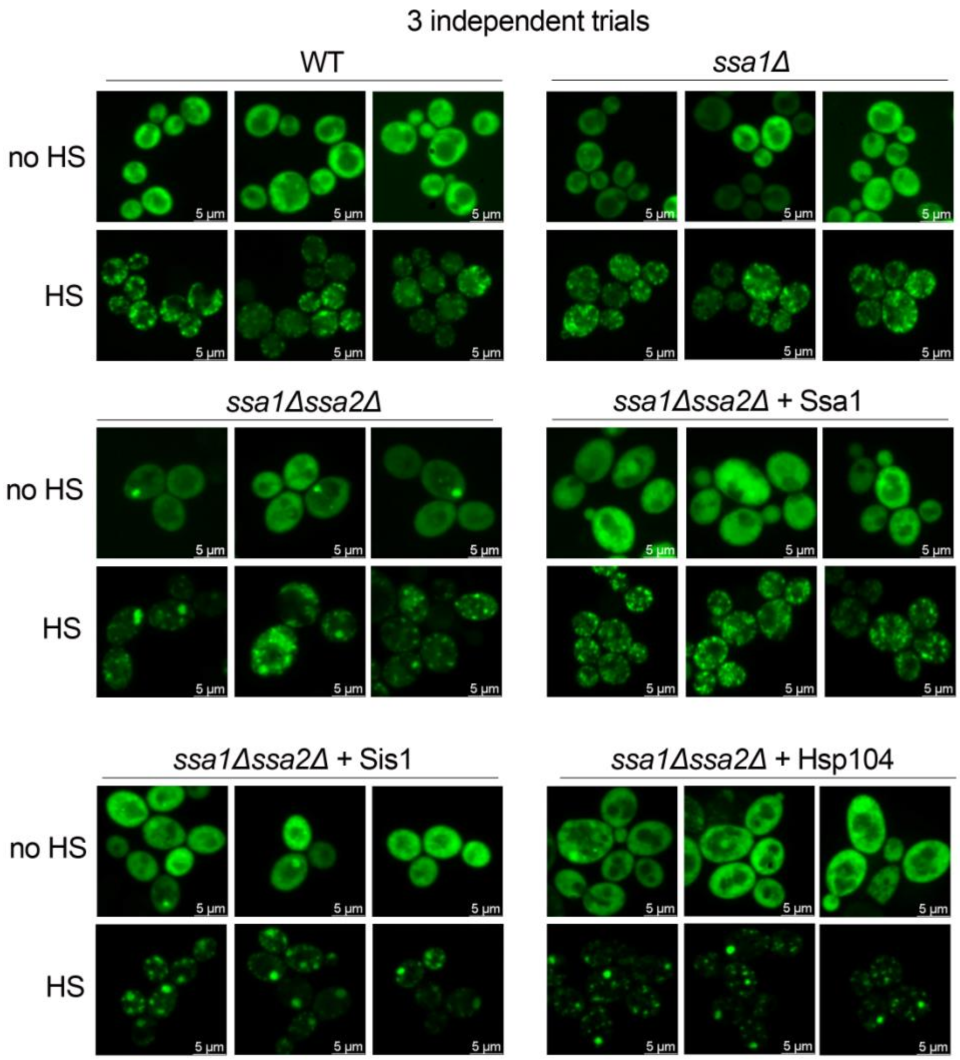


**Supplemental figure 1. Additional representative images relevant to Figure 1A and B.** Three independent images are shown of Pab1-GFP in the indicated strains, without (no HS) and with (HS) heat shock.

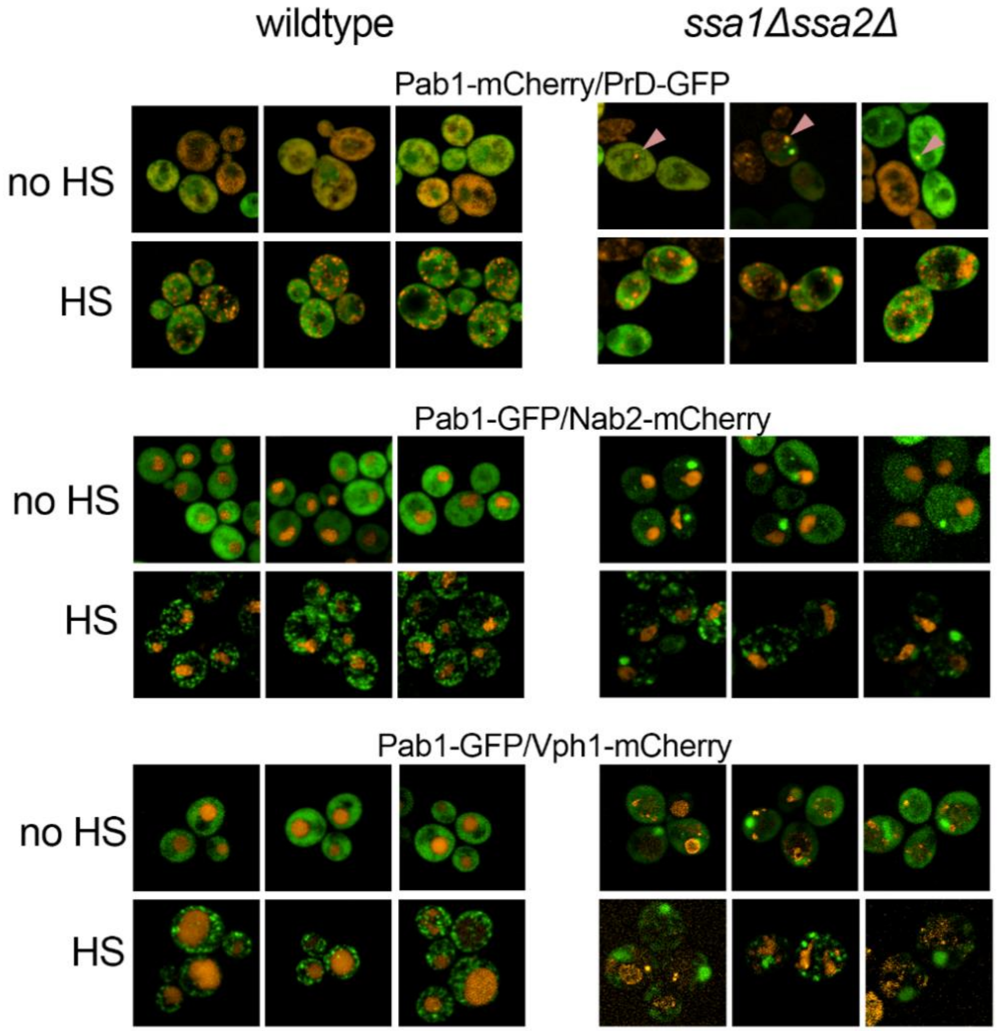


**Supplemental figure 2. Additional merged representative images relevant to Figure 2.** Three independent images are shown of Pab1-GFP in the indicated strains, without (no HS) and with (HS) heat shock. PrD-GFP is the prion domain of Sup35 fused to GFP, Nab2-mCherry localizes to the nucleus, and Vph1-mCherry localizes to the vacuole.

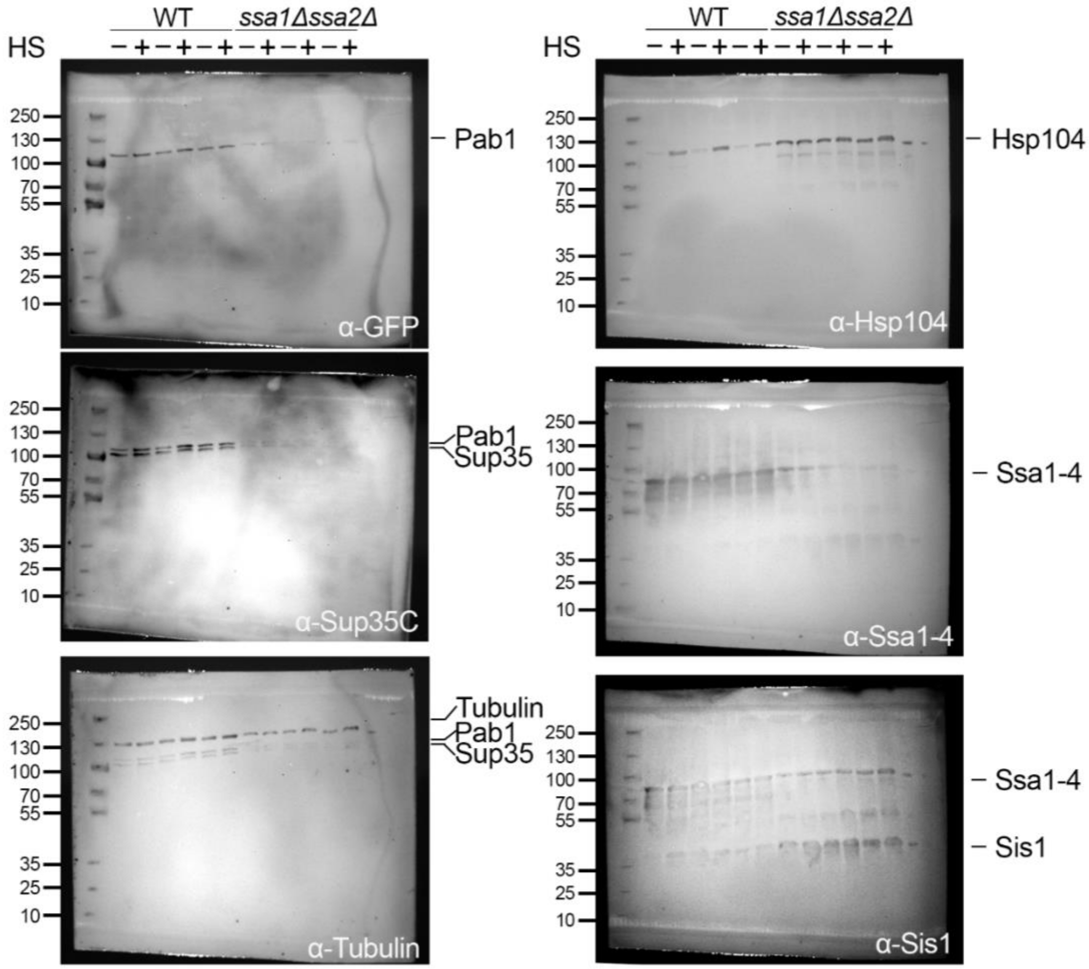


**Supplemental figure 3. Raw Western blot gels relevant to Figure 3.** Raw Western blot images with three samples of wildtype and *ssa1Δssa2Δ* without (-) and with (+) heat shock run on the same gel is shown. Molecular weight markers are shown on the left. Left. The same blot was incubated with antibodies for Pab1 (top), the C-terminus of Sup35 (middle), and Tubulin (bottom) in sequence using the same type of secondary antibody. Protein signals are indicated. Right. Blots from the left were also incubated with antibodies to

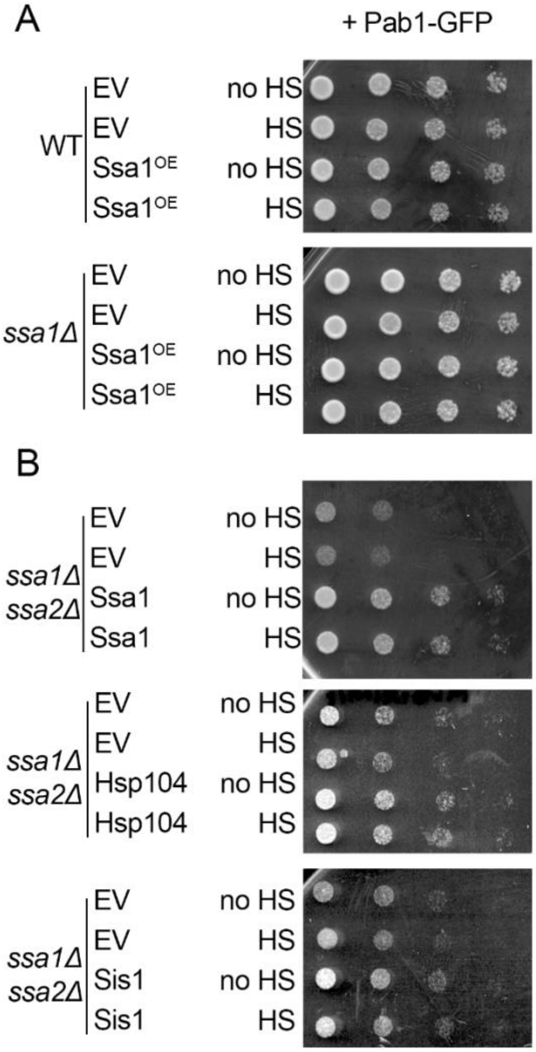


Hsp104 (top), Ssa1-4, (middle), and Sis1 (bottom) using a different secondary antibody. Protein signals are indicated.

**Supplemental figure 4. *ssa1Δssa2Δ* strains exhibit a growth defect that is rescued by Ssa1 overexpression.** A) Wildtype and *ssa1Δ* strains were transformed with Pab1-GFP and either a control plasmid (empty vector; EV) or GPD-Ssa1 overexpression plasmid (Ssa1). Cells were incubated at room temperature (no HS) or at 42°C for 45 minutes (HS). Strains were serially diluted five-fold, spotted on synthetic plasmid-selective media, and grown at 30°C for 2 days. B) Same as A, but in *ssa1Δssa2Δ* containing Pab1-GFP and either EV, GPD-Ssa1, HSE-Hsp104, or GPD-Sis1 overexpression plasmids.

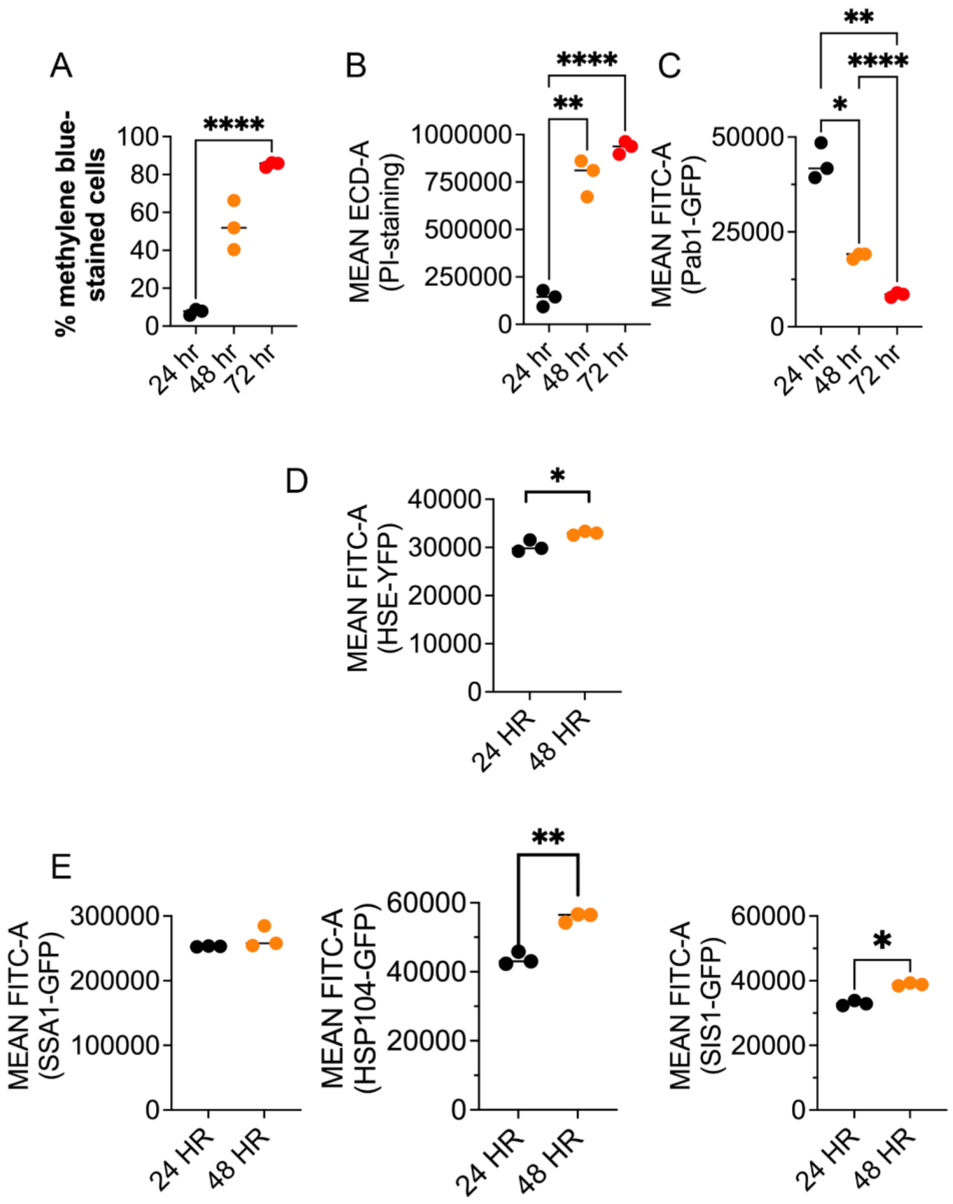


**Supplemental figure 5. Pab1-GFP intensity and cell viability decreases, while HSR and chaperones increase in saturated cultures.** A) Paired cultures were grown for the indicated time and stained with methylene blue to quantify the percentage of dead cells over time via brightfield microscopy. B and C) The same cells in A were also subjected to flow cytometry to quantify the amount of dead cells over time that were stained with propidium iodide (PI) using the ECD-A filter (B) and quantify the mean intensity of Pab1-GFP using the FITC-A filter (C). Means were compared by one way ANOVA (*p=0.0211; **p≤0.0098; ****p≤0.0001) D) Three independent cultures of a WT strain integrated with an HSE-YFP reporter were grown for 48 hours. YFP intensity was quantified through flow cytometry using the FITC-A filter and compared by paired t-test (*p=0.0397). E) Ssa1 (left),

Hsp104 (middle), and Sis1 (right) were endogenously tagged with GFP. Three independent cultures were grown for 48 hours and chaperone-GFP intensity was quantified through flow cytometry using the FITC-A filter and compared by paired t-test (*p=0.0121; **p=0.0075).

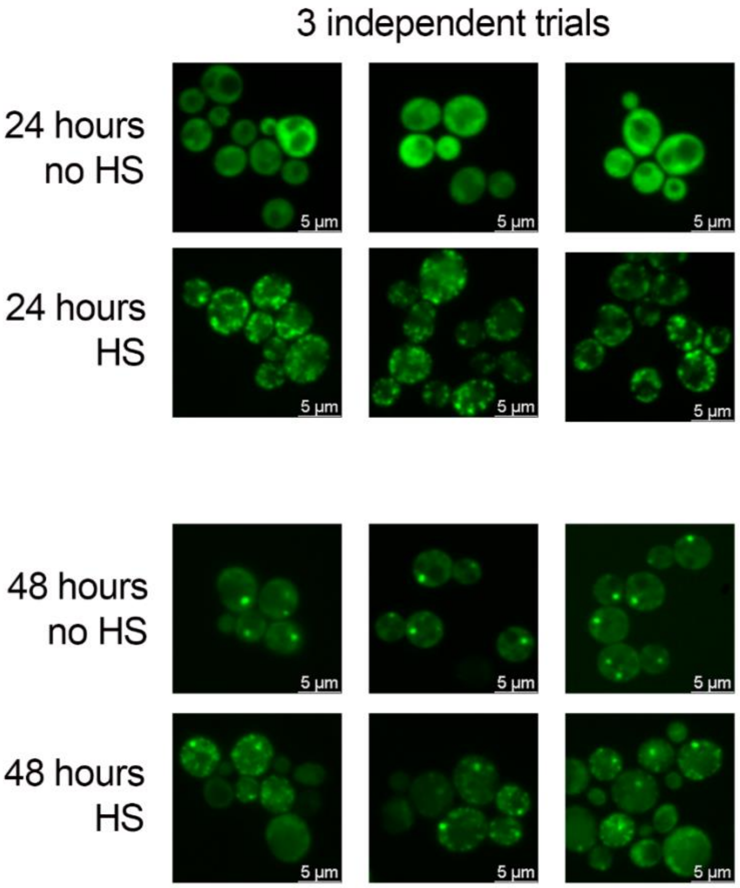


**Supplemental figure 6. Additional representative images relevant to Figure 5.**

**Supplemental table 1. Yeast strains used in this study.**

| Strain # | Genotype | Name | Reference and figure |
|---|---|---|---|
| D230 | Mat**a** ade1-14 ura3-52 leu2-3,112 trp1-289 his3-200 | WT<br>74D-694 | (Chernoff et al., 1995)<br>(All figures) |
| M675 | Mat**a** ade1-14 ura3-52 leu2-3,112 trp1-289 his3-200 ssa1::HIS3 ssa2::NatR | *ssa1Δssa2Δ*<br>74D-694 | This study.<br>Figures 1-4, SF1-4 |
| M652 | Mat**a** ade1-14 ura3-52 leu2-3,112 trp1-289 his3-200 ssa1::HIS3 | *ssa1Δ*<br>74D-694 | This study.<br>Figures 1, 4, SF1, SF4 |
| M761 | Mat**a** ade1-14 ura3-52 leu2-3,112 trp1-289 his3-200<br>Nab2-mCherry::TRP1 | WT<br>74D-694<br>Nab2-mCherry | This study.<br>Figure 2 |
| M757 | Mat**a** ade1-14 ura3-52 leu2-3,112 trp1-289 his3-200 ssa1::HIS3 ssa2::NatR Nab2-mCherry::TRP1 | *ssa1Δssa2Δ*<br>74D-694<br>Nab2-mCherry | This study.<br>Figure 2 |
| M762 | Mat**a** ade1-14 ura3-52 leu2-3,112 trp1-289 his3-200 Vph1-mCherry::TRP1 | WT<br>74D-694<br>Vph1-mCherry | This study.<br>Figure 2 |
| M763 | Mat**a** ade1-14 ura3-52 leu2-3,112 trp1-289 his3-200 ssa1::HIS3 ssa2::NatR<br>Vph1-mCherry::TRP1 | *ssa1Δssa2Δ*<br>74D-694<br>Vph1-mCherry | This study.<br>Figure 2 |

| | | | |
|---|---|---|---|
| M703 | Mat**a** ade1-14 ura3-52 leu2-3,112 trp1-289 his3-200 4xHSE-YFP::LEU2 | WT<br>74D-694<br>4xHSE-YFP | This study.<br>Figure 3, SF5 |
| M707 | Mat**a** ade1-14 ura3-52 leu2-3,112 trp1-289 his3-200 ssa1::HIS3 ssa2::NatR 4xHSE-YFP::LEU2 | *ssa1Δssa2Δ*<br>74D-694<br>4xHSE-YFP | This study.<br>Figure 3 |
| M771 | Mat**a** ade1-14 ura3-52 leu2-3,112 trp1-289 his3-200 ssa1::HIS3 4xHSE-YFP::LEU2 | *ssa1Δ*<br>74D-694<br>4xHSE-YFP | This study.<br>Figure 3 |
| M248 | Matα ade1-14 his3-200 trp1-289 ura3-52 leu2-3,112 HSP104GFP::KANMX6 | Hsp104-GFP<br>74D-694 | Cured of prions<br>(Klaips et al., 2014)<br>SF5 |
| M249 | Mat**a** ade1-14 his3-200 trp1-289 ura3-52 leu2-3,112 SISGFP::KANMX6 | Sis1-GFP<br>74D-694 | Cured of prions<br>(Klaips et al., 2014)<br>SF5 |
| M250 | Mat**a** ade1-14 his3-200 trp1-289 ura3-52 leu2-3,112 SISGFP::KANMX6 | Sis1-GFP<br>74D-694 | Cured of prions<br>(Klaips et al., 2014)<br>SF5 |

**Supplemental table 2. Plasmids used in this study.**

| Plasmid # | Description | Yeast Marker | Reference and figure |
|---|---|---|---|
| 3284 | Pab1-GFP | *URA3* | (Brengues and Parker, 2007) All figures |
| 3285 | Pab1-GFP | *TRP1* | (Brengues and Parker, 2007)Figures 1, 4, SF1, SF4 |
| 3272 | Nab2-mCherry | *TRP1* | (Zhu et al., 2019)Figure 2, SF2 |
| 3273 | Vph1-mCherry | *TRP1* | (Zhu et al., 2019)Figure 2, SF2 |
| 3029 | pCup-Sup35PrD-GFP | *URA3* | (Zhou et al., 2001)Figure 2, SF2 |
| 3384 | Pab1-mCherry | *TRP1* | This study. Figure 2, SF2 |
| 3357 | 4xHSE-YFP | *LEU2* | (Zheng et al., 2016)Figure 3, SF5 |
| 3110 | pRS316 HSE-Hsp104 | *URA3* | (Jackrel and Shorter, 2014)Figures 1, 4, SF1, SF4 |
| 3302 | pAG415 GPD-Ssa1 | *LEU2* | This study. Figures 1, 4, 5, SF1, SF4 |
| 3281 | pAG414 GPD-Sis1 | *TRP1* | This study. Figures 1, 4, SF1, SF4 |
| 3109 | pRS316 HSE-EV | *URA3* | (Jackrel and Shorter, 2014)Figures 1, 4, SF1, SF4 |
| 3168 | pRS315 HSE-EV | *LEU2* | (Jackrel and Shorter, 2014)Figures 1, 4, SF1, SF4 |

**Supplemental table 3. Antibody and conditions used in this study.**

| Antibody | Dilution | Clonality | Vendor |
| --- | --- | --- | --- |
| GFP | 1:5000 | Monoclonal | Roche |
| Sup35C | 1:10000 | Monoclonal | Cocalico Biologicals |
| Ssa1-4 | 1:10000 | Polyclonal | Elizabeth Craig |
| Hsp104 | 1:10000 | Polyclonal | Enzo |
| Sis1 | 1:10000 | Polyclonal | Elizabeth Craig |
| Tubulin | 1:10000 | Monoclonal | Invitrogen |

### REFERENCES


Brengues, M., and R. Parker. 2007. Accumulation of polyadenylated mRNA, Pab1p, eIF4E, and eIF4G with P-bodies in Saccharomyces cerevisiae. *Mol Biol Cell*. 18:2592-2602. PMC1924816

Chernoff, Y.O., S.L. Lindquist, B. Ono, S.G. Inge-Vechtomov, and S.W. Liebman. 1995. Role of the chaperone protein Hsp104 in propagation of the yeast prion-like factor [psi+]. *Science*. 268:880-884. PMID7754373

Jackrel, M.E., and J. Shorter. 2014. Potentiated Hsp104 variants suppress toxicity of diverse neurodegenerative disease-linked proteins. *Dis Model Mech*. 7:1175-1184. PMC4174528

Klaips, C.L., M.L. Hochstrasser, C.R. Langlois, and T.R. Serio. 2014. Spatial quality control bypasses cell-based limitations on proteostasis to promote prion curing. *eLife*. 3. PMC4270096

Zheng, X., J. Krakowiak, N. Patel, A. Beyzavi, J. Ezike, A.S. Khalil, and D. Pincus. 2016. Dynamic control of Hsf1 during heat shock by a chaperone switch and phosphorylation. *eLife*. 5. PMC5127643

Zhou, P., I.L. Derkatch, and S.W. Liebman. 2001. The relationship between visible intracellular aggregates that appear after overexpression of Sup35 and the yeast prion-like elements [PSI(+)] and [PIN(+)]. *Mol Microbiol*. 39:37-46. PMID11123686

Zhu, J., Z.T. Zhang, S.W. Tang, B.S. Zhao, H. Li, J.Z. Song, D. Li, and Z. Xie. 2019. A Validated Set of Fluorescent-Protein-Based Markers for Major Organelles in Yeast (Saccharomyces cerevisiae). *mBio*. 10. PMC6722415
